# Ovule cell wall composition is a maternal determinant of grain size in barley

**DOI:** 10.1101/2022.12.02.518939

**Authors:** Xiujuan Yang, Laura G. Wilkinson, Matthew K. Aubert, Kelly Houston, Neil J. Shirley, Matthew R. Tucker

## Abstract

**Summary:**

- In cereal species, seed and grain size is influenced by growth of the ovule integuments (seed coat), the spikelet hull (lemma and palea) and the filial endosperm. It has remained unclear whether a highly conserved ovule tissue, the nucellus, has any impact on grain size.
- Immunolabelling revealed that the barley nucellus comprises two distinct cell types that differ in terms of cell wall homogalacturonan (HG) accumulation. Transcriptional profiling of the nucellus identified two pectin methylesterase genes, *OVULE PECTIN MODIFIER 1* (*OPM1*) and *OPM2*, which are expressed in the ovule but absent from the seed.
- Ovules from an *opm1 opm2* mutant, and plants expressing an ovule-specific pectin methylesterase inhibitor (PMEI), exhibit reduced HG accumulation. This results in changes to ovule cell size and shape, and ovules that are longer than wild-type controls. At grain maturity, this is manifested as significantly longer grain.
- These findings indicate that cell wall composition during ovule development acts to limit ovule and seed growth. The investigation of ovule PME and PMEI activity reveals an unexpected role of maternal tissues in controlling grain growth prior to fertilisation, one that has been lacking from models exploring improvements in grain size.

## Introduction

Grain size in cereal crops is an important component of yield and quality (Evers & Millar, 2002). Both a seed and a fruit, the grain develops from the female pistil after fertilisation, and its final size is determined by a combination of maternal and paternal inputs. At maturity the grain is comprised of protective tissues (i.e. the maternal hull, pericarp and seed coat) and internal filial seed tissues (i.e. the embryo and endosperm; Li & Li, 2016).

The most abundant filial tissue in the mature grain is the endosperm, the product of fusion between a single sperm and a diploid central cell within the female gametophyte (embryo sac; ES). The timing of endosperm initiation, particularly cellularization, is critical for grain size. This is compromised through mutations in genes from the *HAIKU* (*IKU*) pathway and *Fertilization-Independent Endosperm1* (*FIE1*), which cause precocious cellularization and reduced seed size in Arabidopsis and rice, respectively (Garcia *et al*., 2005; Folsom *et al*., 2014). Final grain size is also influenced by (1) the maternal spikelet hull (i.e. the lemma and palea), which determines the amount of space available for grain expansion; (2) the amount of pericarp tissue, which is positively associated with grain length and weight in wheat (Brinton *et al*., 2017; Herrera & Calderini, 2020); and (3) overall pistil size, which is positively correlated with grain set and weight in several species (Xie *et al*., 2015; Guo *et al*., 2016; Benincasa *et al*., 2017). While many genes and quantitative trait loci (QTL) have been identified as regulators of spikelet size in rice and other cereals (e.g. the ubiquitin– proteasome (UPS) pathway, AP2-transcription factors, G-protein signaling, MAPK signaling and phytohormone signalling; Li *et al*., 2019), relatively little is known about pathways influencing the size of the pistil and its constituent tissues.

In cereal species the upper pistil is composed of the style and stigma to facilitate pollen attachment, while the lower pistil constitutes the ovary, which encloses a single ovule (Fig. S1a). The ovule is a complex structure that represents the progenitor of most seed tissues; it comprises somatic tissues such as the integuments and the nucellus, along with key generative (germline) tissues such as the single ES, which is fertilised to produce the embryo and endosperm (Fig. S1b). After fertilisation, the bulk of the nucellus is rapidly degraded through programmed cell death and the residual cells differentiate into the nucellar projection, which connects to the ovary wall and transfers nutrients into the developing endosperm (Radchuk *et al*., 2011). The nucellar epidermis and two adjoining integuments fuse to become a thin seed coat, which is tightly attached to the ovary wall (pericarp).

The potential impacts of ovule tissue growth, differentiation, and overall size on grain development remain largely unknown. In general, organ size in plants is determined by both cell proliferation and cell expansion (Powell & Lenhard, 2012), which depend on cell cycle and cell wall rigidity, respectively. Cell wall rigidity varies between cells and is strongly influenced by polysaccharide structure and composition (Burton *et al*., 2010; Tucker *et al*., 2018). Pectins, for example, are non-cellulosic polysaccharides that reside in the cell wall matrix and contribute to wall flexibility (Parre and Geitmann, 2005; Haas *et al*., 2020). The abundance and structure of pectin is controlled by a number of enzyme classes (Cantarel *et al*., 2009), including pectin methylesterases (PMEs) that catalyse the de-esterification of pectic domain homogalacturonan (meHG). Demethylesterified homogalacturonan (HG) interacts with Ca2+ to form the so-called ‘egg-box’ structure that crosslinks adjoining polysaccharides, enhancing the rigidity of the cell wall (Du *et al*., 2022). The *PME* gene family usually consists of multiple members in the genomes of higher plants, and specific PMEs are responsible for processes such as organ initiation, stem strength, and plant immunity (Pelloux *et al*., 2007; Wu *et al*., 2018; Francoz *et al*., 2019). Although pectin is less abundant in grasses compared to eudicots such as Arabidopsis (Vogel, 2008), it accumulates in distinct regions during development. For example, the asymmetric distribution of HG and meHG in the tip and shank region of the rice pollen tube is important for tip growth (Li *et al*., 2018; Kim *et al*., 2020). In rice, barley and wheat grain, the nucellar epidermis accumulates HG, although the function is unknown (Palmer *et al*., 2015; Betts *et al*., 2017). Moreover, down-regulation of the pectin biosynthesis gene *GAUT4* in rice and switchgrass led to a reduction in HG levels, decreased recalcitrance in biomass processing, and increased growth (Biswal *et al*., 2018).

Here, we show that pectin (HG) epitopes clearly delineate the inner from outer nucellus in the barley ovule. Using tissue-specific transcriptome profiling and CRISPR/Cas9 gene editing, we identify and analyse the function of two *PME* genes expressed in the nucellus; *OVULE PECTIN MODIFIER1* (*OPM1*) and *OPM2*. Both genes encode apoplastic proteins and are predominantly expressed in somatic nucellar cells adjoining the ES and its precursor cells. While ovules from *opm1* and *opm2* single mutants show reduced HG labelling in the nucellus compared to WT, *opm1 opm2* double mutants lack HG in the nucellus and exhibit significant changes in ovule cell shape. At anthesis, compared with wild-type, *opm1 opm2* mutants produce longer ovules and ovaries. Mirroring the change in ovule/ovary length, the grain length of the *opm1 opm2* mutant is also increased. Expression of a PME inhibitor gene in the nucellus, driven by the nucellus-specific barley *MADS31* promoter, reproduced the increase in ovule, ovary and grain length. Thus, we demonstrate that the ovule can exert maternal control on grain size by adjusting cell wall composition, and identify an intriguing maternal pathway influencing grain development in plants.

## Materials and Methods

### Plant materials and creation of transgenic plants

Barley (*Hordeum vulgare* L.) cultivar Sloop plants were grown in a glasshouse at The Plant Accelerator, Adelaide, Australia, in a 50:50 cocopeat:clay-loam soil mixture (v/v), under 22°C day / 17°C night conditions. Ovules from cv. Sloop plants were used for the initial cell wall immunolabelling screens. In all other experiments, barley cultivar Golden Promise (GP) was used as wild-type and a donor plant for transformation. GP plants were grown in cocopeat soil, at 15 °C light, 12 °C dark, 16 hours daylight with 70% humidity in controlled environment growth chambers (The Plant Accelerator, Waite Campus, The University of Adelaide, Australia).

An optimized CRISPR-Cas9 system for monocots was used for creating barley mutants (Ma *et al*., 2015). Two targets and protospacer adjacent motifs (PAM) for each *OPM* gene were selected from coding regions. The target sites were sequenced in GP before being cloned into vectors to guarantee accurate recognition and editing. Single guide RNA (sgRNA) including target sequences were driven by rice promoters *OsU6a, OsU6b, OsU6c* and *OsU3*, respectively. The sgRNA expression cassettes were amplified using Phusion High-Fidelity DNA polymerase (NEB) and cloned into a binary vector pYLCRISPR/Cas9P_ubi_-H using *Bsa*I site as described (Ma *et al*., 2015). To ectopically express PMEI in the ovule, the *proMADS31::PMEI-eGFP* construct was created by inserting the 2,418 bp *MADS31* (HORVU.MOREX.r3.2HG0190700) promoter and a full length *PMEI* (HORVU.MOREX.r3.1HG0050140) cDNA fused with *eGFP* (enhanced green fluorescent protein) between the *Hind*III and *BstE*II sites of pCAMBIA1301, using the In-Fusion cloning kit (Clontech).

All vectors were transformed into GP using an *Agrobacterium tumefaciens* mediated method as described previously (Harwood *et al*., 2009). Vectors were transformed into the AGL1 strain and overnight cell culture was used to inoculate immature barley embryos. Transgenic seedlings were regenerated from selective media, transferred to cocopeat soil and genotyped by the Phire Plant Direct PCR Kit (Thermo Scientific) and Sanger sequencing. All primers are listed in Table S1.

Between 10-21 primary transgenic lines were generated for each construct and these were confirmed by PCR. In the case of *proMADS31::PMEI-eGFP*, three lines showing GFP expression were chosen for detailed analysis. To confirm the presence of CRISPR-induced edits in *OPM1* and *OPM2*, DNA was extracted from T0 plants and amplified using primers flanking the target sites. Sanger sequencing was used to confirm the presence of edits. Heritability of edits was confirmed in the T1 and T2 generations, and lines were identified that lack the Cas9 construct by PCR. Three independent mutants were identified for each *opm1* and *opm2* single mutant that showed very similar phenotypes in pre-screening. The majority of data included in this study is derived from the strongest independent alleles of each single and double mutant combination.

### Phenotypic analysis and microscopy

For all phenotypic comparisons of pistils, spikelets, ovules and grain, GP plants were grown in a controlled environment growth chamber (see above). Spikelets at anthesis and mature grains were photographed with a Nikon D90 digital camera. The spikelets and grains were collected from the middle region of spikes. Grain and spikelet (awn and sterile florets removed) size was measured using the GrainScan software (Whan *et al*., 2014) by processing a scanned image of grains by Canon CanoScan LiDE 220.

To examine the wholemount details of ovule development by microscopy, central florets were dissected from spikelets located in the middle of the spike just before anthesis, and immediately fixed in ice cold FAA (Formaldehyde Alcohol Acetic Acid, 10%:50%:5% + 35% water). Pistils were dissected and dehydrated in a series of 70, 80, 90, and 100% ethanol and cleared in Hoyer’s solution for two weeks as described previously (Wilkinson & Tucker, 2017). Pistils were imaged using a Zeiss AxioImager M2 with differential contrast microscopy (DIC). Dimensions of ovaries, ovules and embryo sacs were measured using Zeiss ZEN 2012 (Blue) software. Ovules of *proMADS31::PMEI-eGFP* transgenic lines were carefully dissected from pistils and imaged by a Zeiss AxioImager M2 for GFP signals (excitation, 450– 490 nm; emission, 500–550 nm).

### Immunohistochemical assays

To examine the development of ovule cells and tissues by thin sectioning, pistils or whole spikelets were collected and fixed in glass vials containing FAA, dehydrated in a series of 70, 80, 90, and 100% ethanol, and embedded in Technovit 7100 resin (Kulzer Technique) as described by the manufacturer. Replicate pistils were selected at random for sectioning at 1.5 μm on a Leica Ultramicrotome. Immunohistochemical assays were performed as previously described (Betts *et al*., 2017). Mouse antibodies BG1 and anti-callose (1: 100 dilution, Biosupplies, Australia), and rat antibodies JIM13, LM19 and LM20 (1:100 dilution, Kerafast, USA) were used as the primary antibodies to detect (1,3;1,4)-β-glucan, callose, arabinogalactan protein (AGP), de-esterified (HG) and esterified (meHG) pectin, respectively, followed by Alexa Fluor 555 goat anti-mouse/rat IgG (1:200 dilution, Invitrogen) as the secondary antibody. Sections were incubated in Calcofluor White Stain (Sigma) for 1 min for general cell wall staining. After three rinses with water, sections were mounted with 90% glycerol and imaged by a Zeiss AxioImager M2 (for pectin, excitation, 538–562 nm; emission, 570–640 nm; for calcofluor stain, excitation, 335–383 nm; emission 420–470 nm). Sections were rinsed with water again, stained in 0.5% Toluidine blue (w/v) and imaged by Nikon Ni-E optical microscope. The immunohistochemical experiments were carried out at least three times, using at least three randomly selected pistils per stage and genotype.

### Laser microdissection and transcriptome analysis

Whole flowers were collected at approximately stages 8, 8.75 and 9.5 on the Waddington Scale, corresponding to ovule stages Ov4, Ov5, and Ov7, which describe ovules at FG2-4, FG8, and anthesis (Wilkinson, 2019). Similarly, grain samples were collected at 7, 9, 13 and 25 days post anthesis (DPA) (Aubert, 2018). For laser microdissection, the protocol described previously was followed (Okada *et al*., 2013). In brief, samples were fixed in an ice-cold mixture of 3:1 ethanol:acetic acid with 1 mM 1,4-Dithiothreitol (DTT). Samples were stored at 4°C overnight, then transferred into 70% ethanol and stored at -20°C. After dehydration, samples were infiltrated with BMM resin (composed of 40 mL N-butyl methacrylate, 10mL methyl methacrylate, 250 mg benzoin methyl ether (ProSciTech, Australia), and 1mM DTT), placed in BEEM capsules (ProSciTech, Australia) and polymerised under UV light at -20°C for five days. Embedded tissue was serially sectioned in a transverse aspect at 3μm using a Leica UM6 Ultramicrotome (Leica microsystems, Germany) with a glass knife. Sections were placed onto DEPC-treated water droplets on PEN-membrane glass slides (Thermo Fisher Scientific, USA) and evaporated using a 36°C slide warmer. Prior to laser capture, BMM resin was removed from tissue sections by gently rinsing slides in acetone (Sigma) for 10 minutes. Various tissues were collected using a Leica LMD Laser Dissection Microscope (Leica microsystems, Germany) at the Waite Adelaide Microscopy Facility.

Total RNA was extracted from each laser-dissected tissue sample using the Picopure RNA isolation kit (Molecular Devices, USA), with DNase I treatment. Concentration and integrity of the resulting RNA was assessed by a NanoDrop One (Thermo Fisher Scientific, USA). The total RNA was amplified twice with the MessageAmp II aRNA Amplification kit (Thermo Fisher Scientific, USA).

All RNA samples were sent to the Australian Genome Research Facility for sequencing on the Illumina Hiseq platform. Reads were assembled using the barley reference genome (Mascher *et al*., 2017) with CLC Genomics (Qiagen, Netherlands). Normalised read counts (transcripts per million, TPM) were used to reflect the abundance of each gene in each sample. Gene names are annotated as HORVUs, as per the International Barley Sequencing Consortium (IBSC 2012) and IPK (Mayer *et al*., 2012; Mascher *et al*., 2017).

### Expression heat map

TPM values of 35 barley *PME* genes were examined in RNA-sequencing data derived from laser capture dissected ovule and grain samples (Wilkinson, 2019; Aubert, 2018). For each gene, TPM alues were normalized to percentages of the maximum value. All 35 genes were ranked by their maximum expression value. TPM of genes encoding pectin acetylesterases and pectin lyases were also extracted and normalized. The expression heatmap was created using ClustVis (Metsalu & Vilo, 2015).

### RNA extraction and qRT-PCR

Total RNA was extracted from whole spikelets (stages Ov1–Ov5/6), whole pistils (stages Ov7/8–Ov10), floral organs of mature spikelets, seedlings, leaves and stems of Golden Promise using the Spectrum Plant Total RNA Kit (Sigma). 2 μg of RNA was treated by the TURBO DNA-free Kit (Invitrogen) to remove genomic DNA. First strand cDNA was synthesized by 400 U of SuperScriptIII reverse transcriptase (Invitrogen) with oligo-dT as primer. Diluted cDNA was added to iTaq Universal SYBR Green Supermix (BIO-RAD) for real-time quantitative PCR using a QuantStudio Flex 6 (Life Technologies) machine. *HvACTIN7* (HORVU.MOREX.r3.1HG0001540) was used as housekeeping gene for normalization (Li *et al*., 2021). All primers used for qRT-PCR are listed in Table S1.

### RNA in situ hybridization

*OPM1-, OPM2-* and *Histone H4-*specific fragments of coding regions were amplified by PCR using primers fused with T7 polymerase promoter. Digoxigenin (DIG)-labeled NTP (Roche) was integrated into antisense and sense probes during *in vitro* transcription by T7 polymerase (ThermoFisher), according to the manufacturer’s instructions. Fresh spikelets were fixed, dehydrated, embedded in paraffin and sectioned with a Leica rotary microtome RM2265 as described previously (Wu & Wagner, 2012). Slides were assembled using an Insitu Pro VSi robot (Intavis) for pre-hybridization washes, hybridization, stringent washes and immunodetection. An antibody conjugate anti-DIG-AP (Roche, 1:1000 dilution) and NBT/BCIP substrate were used to visualize hybridization results. All primers used are listed in Table S1.

### Subcellular localization

*35Spro::OPM1-eGFP* and *35Spro::OPM2-eGFP* were created by inserting full length coding sequence of *OPM1* and *OPM2* fused with eGFP into *pCAMBIA1301* using *Nco*I and *BstE*II sites. Plasmids were transformed into onion (*Allium cepa*) epidermal cells using a particle delivery system (BIO-RAD) as previously described (Yang *et al*., 2018). Fluorescence images were taken from an A1R Laser Scanning Confocal Microscope (Nikon) (for GFP, excitation, 488 nm; emission, 505-520 nm). Images were processed with NIS-Elements Viewer 4.20 (Nikon). All primers used are listed in Table S1.

### Phylogenetic analysis

Phylogenetic trees were generated using alignments of full-length amino acid sequences of all PMEs in the barley genome and homologs of OPM1 and OPM2 from other higher plant species. Neighbour-joining methods (MEGA version 7.0) were used with a p-distance model and pairwise deletion and bootstrap. The maximum parsimony method of MEGA also was used to support the neighbour-joining tree using the default parameter.

### Haplotype analysis

Using published exome capture data (Russell *et al*., 2016; Mascher *et al*., 2017; Bustos-Korts *et al*., 2019), SNPs were examined in 210 georeferenced landrace barley and 109 *H. spontaneum* accessions for *OPM1* and *OPM2*. These polymorphisms were mapped onto the corresponding gene model to identify the location of the SNP within the gene, and to determine if the polymorphism was synonymous or non-synonymous. Haplotypes were constructed and plotted geographically.

### Replication and statistical analysis of tissue dimensions

Samples were collected from plants grown in a random design in a controlled environment chamber (see above). Statistical tests were selected based on simulations and advice from SAGI (Statistics for the Australian Grains Industry), and used to determine whether differences in ovule, pistil, grain and spikelet measurements were significant between replicate plants, samples and/or genotypes.

For histological sectioning (Fig. 3), a pool of 200 pistils were collected from 6 replicate plants (between 30-36 pistils per plant) for each genotype. Eight pistils per genotype were selected at random for sectioning, and three thin sections were analysed for each pistil. Multiple simulations were run based on the number of plants and pistils to determine the effective degrees of freedom. The different positions and zones were analysed using a repeated-measures mutli-response linear model analysis (ANOVA of repeated measures) to assess the effects of genotype, section and section by genotype. The design was balanced, and the residuals of the model satisfied the assumptions of normality and sphericity. Data from the three independent thin sections from the same ovule were collected to account for variation in the exact position of the sagittal plane; however statistical analysis suggested the effects of section or section by genotype were not significant.

For the whole ovule and embryo sac measurements (Fig. 4, Fig. S10), 20 pistils were selected at random from the same pool of 200 pistils (described above). Analysis suggested the ovules and ovaries of the pistils collected were at the same stage of development across the collection, and represent 4-5 replicate plants. A version of ANOVA was used that does not assume homogeneity of variance in residuals. Multivariate analysis identified significant differences between genotypes.

For grain analysis (Fig. 4), between 100-134 grain from 3-4 independent replicate plants were collected for each genotype. A stratified ANOVA was undertaken to assess for significance using the stratum of individual plants within genotypes, and seeds within plants.

For analysis of spikelet dimensions (Fig. S11), 41 spikelets from wild type and 49 spikelets from *opm1 opm2* were collected from at least 3 spikes and 3 replicate plants for each genotype. Data was analysed using an equivalence test and a 5% threshold to assess for differences.

## Results

In mature barley ovules, we observed two types of somatic cells within the nucellus (Fig. 1a). Outer nucellar cells adjacent to the integuments and distal to the ES were small in size, densely packed and contained a prominent nucleus (Fig. 1b). In contrast, approximately four to six layers of inner nucellar cells adjoining the ES were enlarged and highly vacuolated (Fig. 1c). We considered that the morphological differentiation between outer and inner nucellar cells might also reflect differences in cell wall composition. To test which cell wall components were present in the ovule, immunohistochemical assays were used to assess callose, (1,3;1,4)-β-glucan, arabinogalactan protein (AGP), methylesterified homogalacturonan (HG) and demethylesterified HG accumulation in wild-type ovules at two stages of development (Ov3 and Ov10). Although most epitopes were detected, the demethylesterified HG epitopes recognized by the LM19 antibody were particularly abundant throughout the walls of inner nucellar cells (Fig. S2, Fig. 1d, 1f). In contrast, the outer nucellar cells showed limited LM19 labelling that was restricted to the junctions between cells (Fig. 1e), suggesting a low level of HG. The small size of cells in the outer nucellus also implies active cell divisions. mRNA *in situ* hybridization of barley *Histone H4* was used as a marker for mitosis, identifying cells in S phase of the cell cycle (Li *et al*., 2021; Fig. 1g). Complementing the spatial arrangement of outer and inner nucellus at ovule maturity, cell division activities were predominantly detected in the outer nucellus (Fig. 1h), while the inner nucellus cells were apparently not dividing (Fig. 1i). The differentiation between outer and inner nucellus in terms of cell shape, cell wall components and cell division activity suggest these two areas of nucellus may contribute to ovule growth in different ways, via cell division in the former and cell expansion in the latter.

**Fig. 1:**
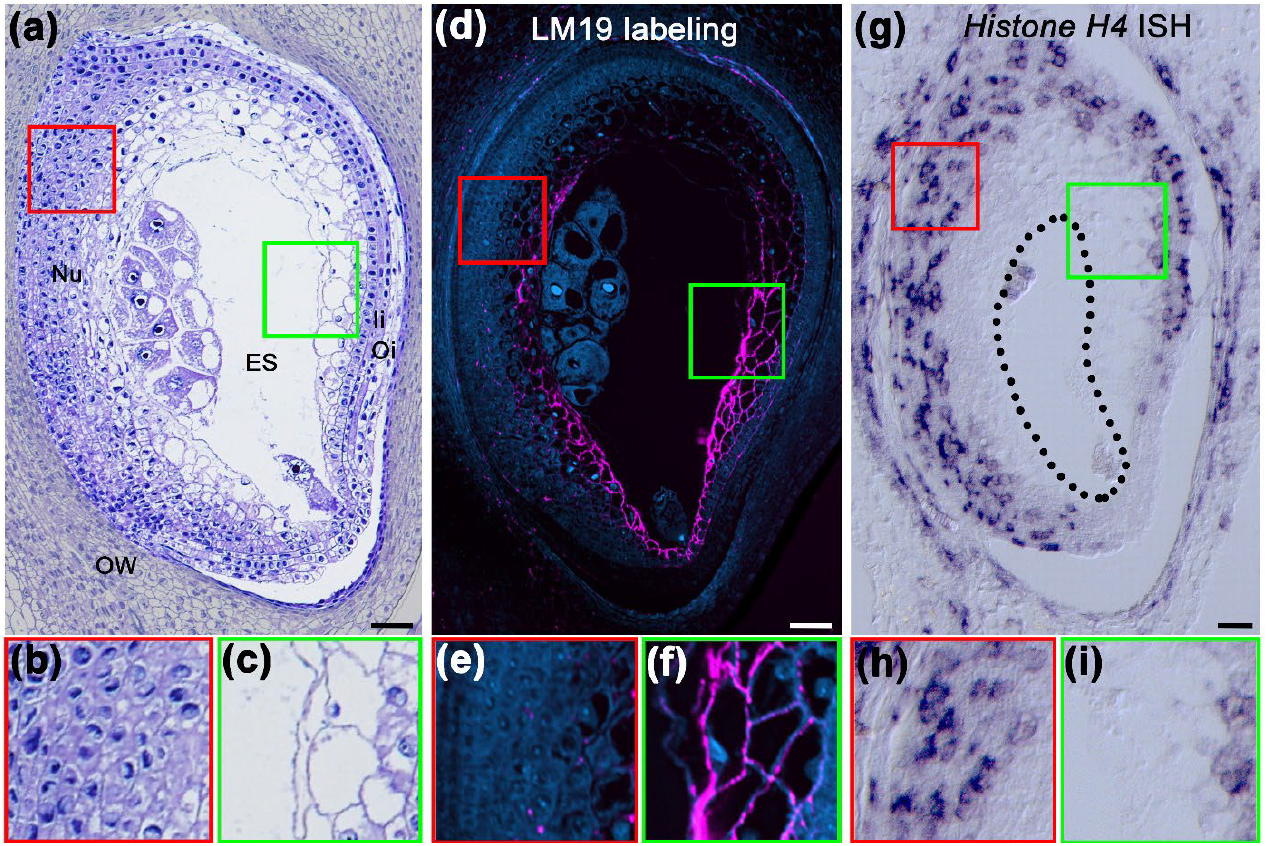
Differentiation of inner and outer nucellus cells in the barley ovule. **(a)** Longitudinal section of a barley ovule at anthesis (ovule stage 10; Ov10). OW, ovary wall; Oi, outer integument; Ii, inner integument; ES, embryo sac. **(b)** and **(c)**, magnifications for outer (red crop) and inner (green crop) nucellus, respectively. **(d)** Immunohistochemical assay of de-esterified pectin (antibody LM19) showing localization in a barley ovule at the same stage with (a). The inner nucellus was intensively labeled by LM19 (shown in magenta). **(e)** and **(f)**, magnifications for outer (red crop) and inner (green crop) nucellus, respectively. **(g)** *Histone H4* mRNA *in situ* hybridization in a barley ovule at the same stage as (a) and (b). Cell division activity was detected in outer nucellus, integuments and ovary wall. **(h)** and **(i)**, magnifications for outer (red crop) and inner (green crop) nucellus, respectively. Bars = 50 μm.

To investigate how the distinct pectin pattern in the nucellus is established and affects ovule development, we generated a laser micro-dissected tissue-specific transcriptome resource for the barley ovule, ovary and grain, and searched for putative ovule-expressed *PME* genes. Among the 42 annotated *PME* genes in the barley genome, 35 were expressed in various tissues of the ovule, ovary and/or grain (Fig. 2a). Two candidates, HORVU.MOREX.r3.2HG0213860 and HORVU.MOREX.r3.6HG0622780, were identified as being ovule-enriched and were chosen for further analysis over grain-expressed *PMEs*, low expressed *PMEs*, and those that were not expressed in the nucellus (Fig. 2a and Fig. S1a). These genes were named as *OVULE PECTIN MODIFIER 1* (*OPM1*; composed of HORVU.MOREX.r3.2HG0213860/90; Figs. S3a and S5a) and *OPM2*, respectively. Quantitative reverse transcription PCR (qRT-PCR) analysis confirmed that *OPM1* and *OPM2* were highly expressed in the pistil compared with other floral organs or vegetative tissues (Fig. 2b). On a temporal scale, *OPM1* and *OPM2* initiated expression as the ES began dividing (Stage Ov7/8; Wilkinson *et al*., 2019) and increased in abundance as the ovule approached maturity (Fig. 2b).

**Fig. 2:**
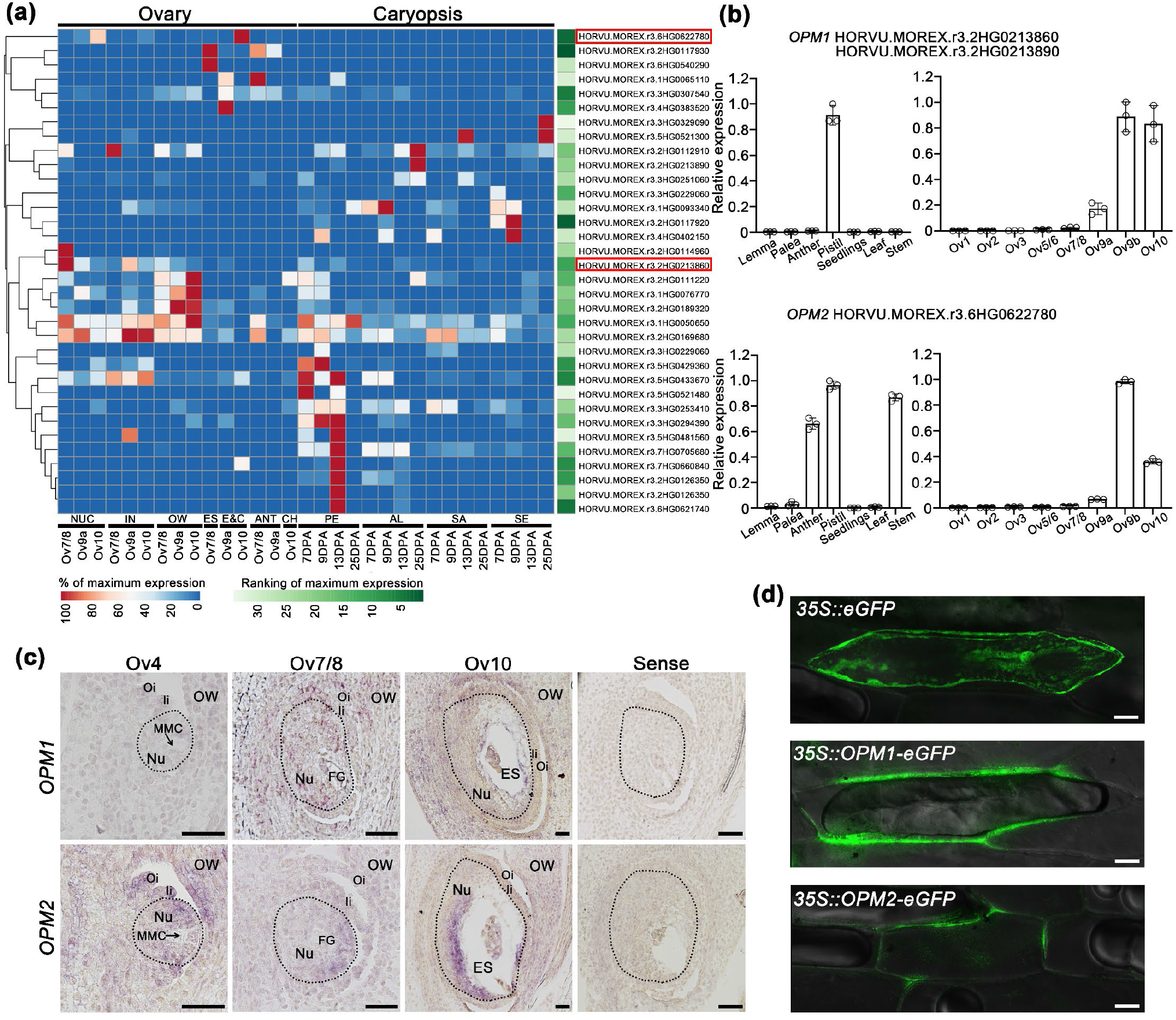
Molecular characteristics of *OPM1* and *OPM2*. **(a)** Heatmap visualization of the expression levels of *PME* genes in the ovule and caryopsis. Transcript abundance values normalized to 1. 35 *PME* genes were ranked according to the maximum expression levels, as shown by the green bar. Ov7/8, female gametophyte mitosis; Ov9, female gametophyte maturity; Ov10, female gametophyte anthesis. DPA, days post anthesis. NUC, nucellus; IN, integuments; OW, ovary wall; ES, embryo sac; E&C, egg cell and central cell; ANT, antipodal cells; CH, chalaza; PE, pericarp; AL, aleurone; SA, sub-aleurone; SE, starchy endosperm. **(b)** qRT-PCR showing relative expression levels of *OPM1* and *OPM2* in various floral organs (left) and various stages of ovule development (right). Error bars represent SD (n=3). *HvACTIN7* serves as reference gene (Li *et al*., 2021). Ov1, ovule primordium; Ov2, archesporial cell; Ov3, megaspore mother cell; Ov5/6, functional megaspore; Ov7/8, female gametophyte mitosis; Ov9a, mature female gametophyte; Ov9b, female gametophyte expansion; Ov10, female gametophyte anthesis. **(c)** *In situ* hybridization of *OPM1* and *OPM2* in wild-type ovules. Sense probes were used as control. MMC, megaspore mother cell; Nu, nucellus; OW, ovary wall; Oi, outer integument; Ii, inner integument; FG, female gametophyte; ES, embryo sac. The black dashed lines indicate the nucellus. Bars = 50 μm. **(d)** Subcellular localizations of OPM1-eGFP and OPM2-eGFP in onion epidermal cells. EGFP was expressed as a control. Bars = 50 μm.

The laser microdissection results were confirmed by mRNA *in situ* hybridization, which showed that *OPM1* mRNA was detected in the nucellus, integuments and parts of the ovary wall that adjoin the nucellus during ES development. At stage Ov10 (anthesis), *OPM1* expression was mainly localised to the innermost layers of inner nucellus (Fig. 2c). *OPM2* was detected in the young nucellus when germline meiosis was initiating (stage Ov4), particularly in cells directly adjoining the germline. This pattern continued in subsequent stages, especially in the mature ovule, when the spatial pattern of *OPM2* expression matched the inner nucellus domain labelled by LM19 (compare Figs. 2c and 1d).

It is widely accepted that pectins are secreted from the Golgi to the cell wall in an esterified form (meHG), where they are modified by PME proteins (Micheli, 2001; reviewed in Du *et al*., 2022). To investigate subcellular localization of OPM1 and OPM2, we transiently expressed both proteins fused to enhanced green fluorescence protein (eGFP) in onion epidermal cells. Compared with eGFP driven by the CaMV 35S promoter, OPM1-eGFP and OPM2-eGFP fusions showed obvious apoplastic localization, suggesting OPM1 and OPM2 are secretory proteins that function in the cell wall (Fig. 2d). Supporting this, TargetP 2.0 analysis predicted the existence of signal peptides and non-organelle localization for both proteins (Fig.S3b).

Although both proteins accumulate in the apoplast and both mRNAs show overlapping spatial and temporal expression in the ovule and pistil, phylogenetic analysis showed that OPM1 and OPM2 are not close homologues in barley (Fig. S4a). Moreover, while OPM2 is relatively conserved in cereal crops, OPM1 appears to be restricted to the Triticeae tribe (Fig. S4b). We therefore considered that the two proteins might reflect distinct origins, but share similar PME activity in ovule tissues during development.

To assess the potential role of these genes in pectin modification during ovule development, we created single and double mutants of *OPM1* and *OPM2* using an optimized CRISPR/Cas9 genome editing system that allows for multiple guide RNAs and targets in one vector (Ma *et al*., 2015). The resultant editing events included insertions and deletions within one or both target genes, and a unique deletion of a large fragment between the two gRNA targets for OPM1 (Fig. S5a). Most INDELs resulted in frameshift mutations and/or premature stop codons in the amino acid sequence of each individual gene, and in several cases, both *OPM1* and *OPM2* (Fig. S5b).

To assess the effect of the mutations, we examined the distribution of HG in wild-type Golden Promise, putative null single mutants, and *opm1 opm2* double mutant ovules. In wild-type, LM19-labeled HG accumulated in the cell walls of nucellar cells directly surrounding the dividing ES (Fig. 3a). Compared to other nucellar cells that were small and almost rounded, these labelled cells were polygonal and had straight edges (Fig. 3d) and were inactive in cell division, as shown by *in situ* hybridization of *Histone H4* (Fig. 3g). During expansion and maturation of the ES in wild-type ovules (Ov9b and Ov10 stages), the nucellar cells close to the ES maintained a high level of HG (Fig. 3b, c), exhibited regular shape (Fig. 3e, f) and did not express a cell division marker (Fig. 3h, i). The nucellar cells of the *opm1* and *opm2* single mutants were only partially labelled by LM19. Compared to wild-type, LM19 labelling was generally reduced in *opm2*, while the labelled region was smaller in *opm1* and decreased further over time (Fig. S6b). In *opm1 opm2* double mutants, the loss of *PME* function dramatically decreased the abundance of de-esterified HG in both the inner and outer nucellus, since LM19-labelling was not detected in *opm1 opm2* ovules during ES mitosis or at anthesis (Fig. 3j, l; Fig. S6a). Hence, both *OPM1* and *OPM2* contribute to the modification of pectin structure in the barley ovule.

**Fig. 3:**
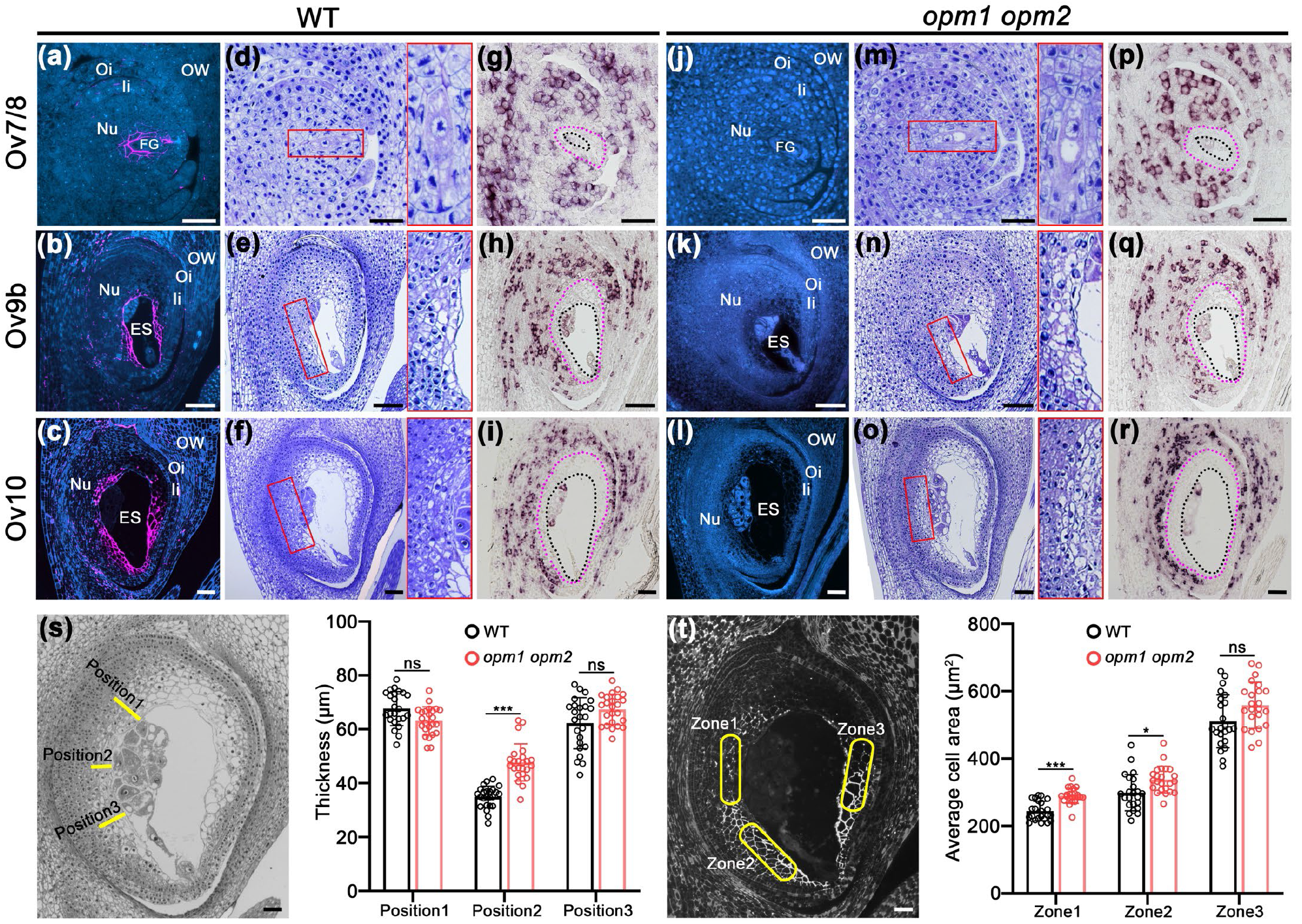
*OPM1* and *OPM2* contribute to pectin de-esterification in the barley ovule.(a–c) Immunohistochemical assay of de-esterified pectin (detected by LM19 antibody) in wild-type ovules at different stages. Ov7/8, female gametophyte mitosis; Ov9b, female gametophyte expansion; Ov10, female gametophyte anthesis. **(d–f)** Cytological analysis of wild-type ovules at the same stages as (a–c). Magnified views of the inner nucellar cells are shown in red rectangles. **(g–i)** *Histone H4 in situ* hybridization in wild-type ovules at the same stage as (a–c). The black outline marks the embryo sac and pink outline marks the inner nucellus region. **(j–l)** Immunohistochemical assay of de-esterified pectin (detected by LM19 antibody) in *opm1 opm2* ovules at different stages. **(m–o)** Cytological analysis of *opm1 opm2* ovules at the same stages as (j–l). Magnified views of the inner nucellar cells are shown in in red rectangles. **(p–r)** *Histone H4 in situ* hybridization in *opm1 opm2* ovules at the same stage as (j–l). **(s)** Thickness of inner nucellus adjacent to the antipodal cell group. Three positions were examined and are indicated. All error bars represent SD, *n =* 8 (3 sections × 8 pistils from a pool of ∼200 pistils from 6 plants). ANOVA test for repeated measures. ***, *P* ≤0.001; ns, not significant. **(t)** Average cell area of inner nucellar cells that are significantly labelled by LM19. Three zones were examined and are indicated. All error bars represent SD, *n =* 8 (3 sections × 8 pistils from a pool of ∼200 pistils from 6 plants per genotype). ANOVA test for repeated measures. ***, *P* ≤0.001; *, *P* ≤ 0.05; ns, not significant. OW, ovary wall; Oi, outer integument; Ii, inner integument; FG, female gametophyte; ES, embryo sac. Bars = 50 μm.

To confirm the role of PME activity in the barley ovule, we also created a transgenic line expressing a PME inhibitor (PMEI) fused to eGFP and driven by the promoter of the ovule-specific *MADS31* gene. Activity of this promoter overlaps with ovule cells that accumulate *OPM1* and *OPM2* transcript, confirmed by strong expression of eGFP in the ovule (Fig. S7a). The degree of LM19 labelling was heavily reduced in the ovules of transgenic lines where PMEI-eGFP was abundantly expressed (as indicated by eGFP signal), similar to *opm1 opm2*, indicating the PMEI was able to partially neutralize PME in a dominant manner (Fig. S7b–d).

Morphologically, the inner nucellar cells in *opm1 opm2* mutants were irregular and distorted, with either a more rounded or compressed shape compared with the corresponding cells in wild-type (Fig. 3m–o). In addition, the inner nucellus in *opm1 opm2* appeared expanded, especially in the area adjoining the antipodal cells (Fig. 3o, r). This did not appear to correlate with any change in cell division, since nucellar cells in *opm1 opm2* mutants showed a similar pattern of *H4* expression compared to wild-type (Fig. 3p–r). To quantify these differences, the inner nucellus thickness was measured at three positions using the antipodal cells as a positional reference (Fig. 3). Thin sections were collected from eight pistils for each genotype, and measured at three distinct positions (1, 2 and 3) in each ovule (Fig. 3s). Consistent with the cytological observations (Fig. 3o, r), the inner nucellus thickness was significantly increased in *opm1 opm2* mutants at position 2, compared with wild-type ovules (Fig. 3s). Moreover, three zones that were heavily labelled by LM19 were examined to determine if loss of LM19-labelled HG might influence cell expansion. In *opm1 opm2* ovules, nucellar cells in several zones had significantly larger average cell areas, compared with wild-type ovules (Fig. 3t). This suggests that loss of de-esterified HG may lead to cell expansion, but in a spatially-restricted manner dependent on constraints from neighbouring tissues.

Deactivation of PMEs might be expected to result in the accumulation of meHG in cell walls. To test this, we utilized the LM20 antibody to examine the abundance of meHG in both wild-type and *opm1 opm2* ovules. Surprisingly, only few punctate signals were present in the wild-type nucellus during mid to late stages of ovule development (Fig. S8a–c), and there was no increase in labelling detected in *opm1 opm2* (Fig. S8e–g). As an internal control, LM20 was shown to label meHG in the anther and ovary cell walls (Fig. S4e, h), suggesting that meHG in *opm1 opm2* ovule may be rapidly degraded or recycled from the cell walls. Supporting this, multiple pectate lyases and polygalacturonases were found to be expressed in the barley ovule (Fig. S9). Several of these genes that show expression in the nucellus might participate in pectin degradation.

Considering the nucellus occupies a large proportion of the whole ovule (Wilkinson *et al*., 2019), we investigated if changes in HG accumulation due to reduced PME activity affected ovule size, ovary size and grain size. In both the *opm1 opm2* mutant and *pMADS31*::*PMEI-eGFP* transgenic plants, ovule length was increased while ovule width remained similar compared to wild-type (Fig. 4a–c). The ES was wider but not longer in the *opm1 opm2* mutant and *pMADS31*::*PMEI-eGFP* plants (Figs. S10a–c), indicating the changes in ovule length are driven mainly by the nucellus. In terms of ovary size, both ovary length and width were increased in the *opm1 opm2* mutant and *pMADS31*::*PMEI-eGFP* lines (Fig. 4d–f). Ovary length and width were positively correlated (Fig. S10d), while the correlation between ovule length and ovary length was stronger than that of ovule width and ovary width (Figs. S10e, f). Taken together, this suggests that longitudinal growth of the ovule is likely to be a key determinant of ovary length, which then has a secondary effect on ovary width.

**Fig. 4:**
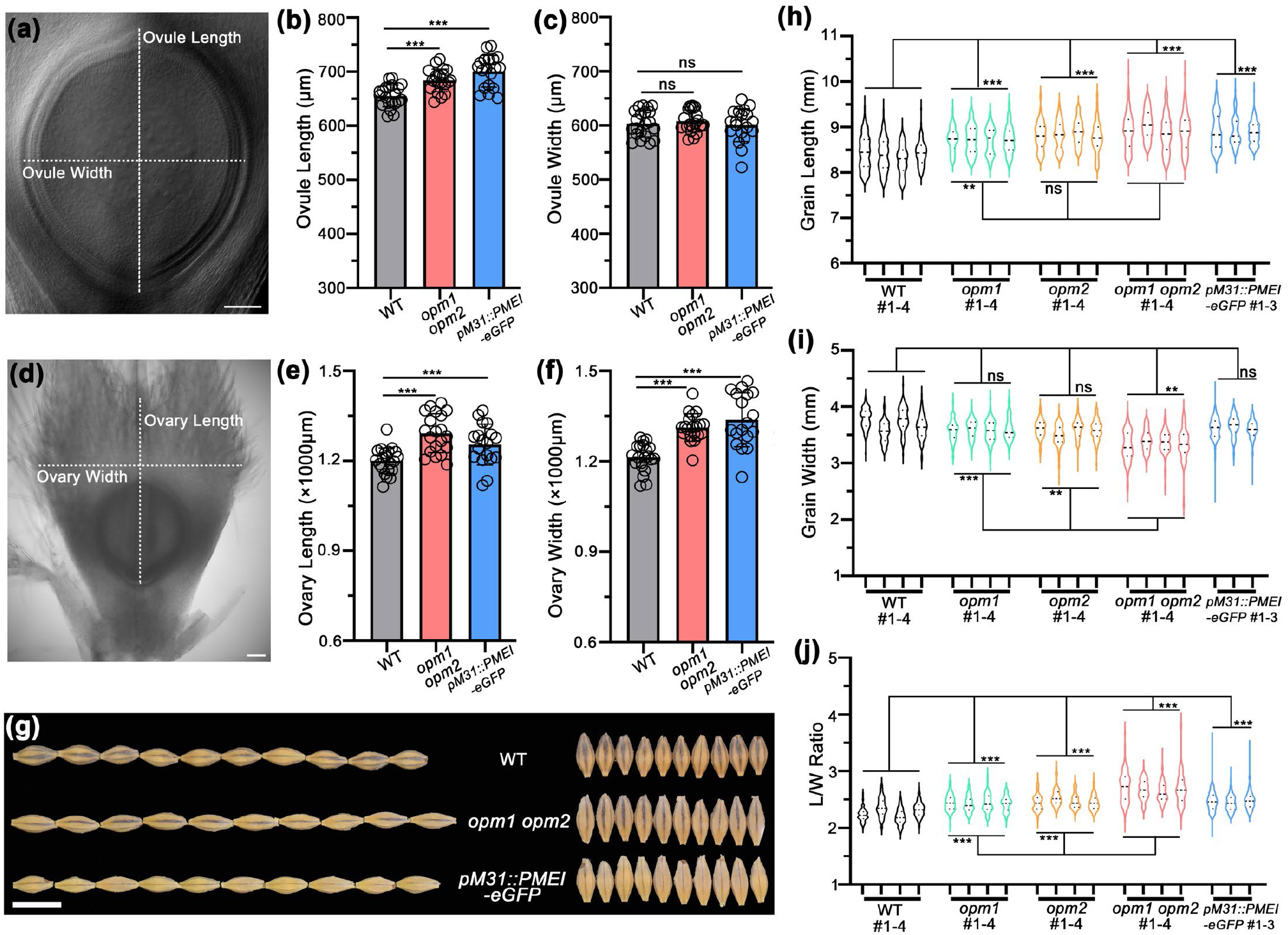
OPM1 and OPM2 affect ovule, ovary and grain growth. **(a)** Ovule dimensions. Ovule length is the distance between the top and the bottom of outer integument. Ovule width is the widest distance between the edges of outer integument. Bar = 100 μm. **(b)** and **(c)** Measurements of ovule dimensions in wild-type, *opm1 opm2* and *pMADS31::PMEI-eGFP* ovules. All error bars represent SD, *n =* 20 pistils (from a pool of ∼200 pistils from 6 plants). ANOVA test. ***, *P* ≤0.001; ns, not significant. **(d)** Ovary dimensions. Ovary length is the distance between the end of style and the bottom of outer integument. Ovary width is the largest transverse diameter of ovary. Bar = 100 μm. **(e)** and **(f)** Measurements of ovary dimensions in wild-type, *opm1 opm2* and *pMADS31::PMEI-eGFP* ovaries. All error bars represent SD, *n =* 20 pistils (from a pool of ∼200 pistils from 6 plants). ANOVA test. ***, *P* ≤0.001. **(g)** Grain shape comparisons of wild-type, *opm1, opm2, opm1 opm2* and *pMADS31::PMEI-eGFP* grains. Bar = 1 cm. **(h–j)** Measurements of grain length (h), grain width (i) and length/width ratio (j) in wild-type, *opm1 opm2, opm1, opm2* and *pMADS31::PMEI-eGFP* grains. *n =* 400 (100–134 grains form 3–4 plants). Stratified ANOVA test. ***, *P* ≤0.001; ** *P* ≤ 0.01; ns, not significant.

We also measured grain size to verify the potential effects of changes in pre-fertilisation ovule growth on grain development. In single *opm1* and *opm2* mutants, *opm1 opm2* double mutants and *pMADS31*::*PMEI-eGFP* lines, grains were longer than those in wild-type (Fig. 4g, h), with no significant change in grain width except for *opm1 opm2* mutants (Fig. 4g, i). As a consequence, the length/width (L/W) ratios of *opm1, opm2, opm1 opm2* and *pMADS31*::*PMEI-eGFP* grains were greater than those of wild-type (Fig. 4j), showing a significant change in grain shape. This affect appears to be specific to the ovule and ovary, since we found no significant change in either length or width of spikelets compared to wild-type (Fig. S11a–c). Hence, the change of grain shape is not caused by differences in lemma and palea growth or spikelet shape.

Collectively, these results demonstrate that changes in organ shape can be passed from the ovule to the ovary, and further to the grain. Since *OPM1* and *OPM2* are predominantly expressed in the nucellus, which is a sporophytic tissue that degrades quickly after fertilisation, *OPM1* and *OPM2* appear to be maternal regulators of grain development in barley.

## Discussion

Organ size is controlled by a range of genetic and environmental factors (Conlon & Raff, 1999). In plants, early stages of organ development are generally defined by cell proliferation, where cell size remains relatively constant, followed by a post-mitotic expansion phase where ploidy may increase, the cell wall is reinforced, and cells become bigger (Powell & Lenhard, 2012). In the context of grain, final size depends on complicated interactions between genetically distinct tissues that restrict or promote growth (Jakobsson & Eriksson, 2000). Diploid maternal tissues such as the seed coat (derived from the maternal ovule integuments) and surrounding hull tissues (lemma and palea), provide physical constraints to limit seed size (reviewed in Li *et al*., 2018). On the other hand, filial components that promote seed size after fertilization include the number of endosperm divisions and the timing of cellularisation (cell wall formation; Wu et al., 2016). Although development of the ovary prior to anthesis also appears to be a determinant of grain growth (Calderini *et al*., 1999; Xie *et al*., 2015; Guo *et al*., 2016), most genetic, physical and nutritional features that affect seed and grain size have been documented to act after fertilization. Relatively little attention has been paid towards understanding the involvement of transient cell types in the ovule that precede the main events of seed development.

Here we show that prior to fertilisation, the bulk of the barley ovule consists of two prominent somatic cell types. The inner nucellus cells adjoining the embryo sac exhibit walls enriched in de-esterified pectin (HG) and low cell division activity, while the outer nucellus comprises cells with a low level of HG and highly active cell division. By generating a tissue-specific expression atlas of all PME genes in the barley ovule and grain, we identified candidates potentially responsible for this difference in composition: *OPM1* and *OPM2*. Removal of *OPM1* and *OPM2* through gene editing essentially eliminated HG epitopes in the barley ovule, both in the inner and outer nucellus, and altered the morphology of inner nucellus cells.

Although cell division remained inactive in the inner nucellus of *opm1 opm2* ovules, cell shape became more rounded and elongated, and as a result, ovules and ovaries of the *opm1 opm2* mutant grow longer and wider. As a developmental outcome, *opm1 opm2* grains exhibit increased length. These findings identify a new link between the composition of pre-anthesis ovule tissues and post-fertilisation cereal grain growth. Whether these genes have contributed to seed size in natural barley populations or breeding lines remains unclear. However, screening of wild and landrace barley panels (Russell *et al*., 2016, Bustos-Korts *et al*., 2019) revealed several single nucleotide polymorphisms (SNPs) in the *OPM1* and *OPM2* genes that result in non-synonymous amino acid changes, and vary depending on geography (Fig. S12). Further studies will provide insight regarding the relationship between these SNPs and functional differences in OPM1 and OPM2, and features of ovule, ovary and grain development.

By comparing the ovules of major cereal crops such as barley, wheat (Chaban *et al*., 2011), rice (Ma *et al*., 2019) and maize (Srilunchang *et al*., 2010), we propose that the inner nucellus “zone” is a common characteristic of cereal ovules; hence the underlying differentiation process might be conserved. Despite this, the region is poorly characterized, and a biological function has remained elusive. In rice ovules, the inner nucellus is enriched in arabinogalactan proteins (AGPs), implying possible functions in signal transduction and tissue patterning (Ma *et al*., 2019). In wheat ovules, enlarged nucellar cells accumulate the pectic polysaccharide rhamnogalacturonan (Chateigner-Boutin et al., 2014). In Arabidopsis, de-esterified HG accumulates at the micropyle to prevent multiple sperm cells from fertilizing the egg cell, and the accumulation of HG is triggered by fertilisation (Duan *et al*., 2020). The pattern of HG distribution we observed in barley ovules was first detected during megaspore mother cell specification, much earlier than fertilisation, and the fertility of the *opm1 opm2* mutant appears normal. This implies a distinct role for this type of pectin during nucellus development in cereal ovules.

HG plays dual roles in regulating cell wall elasticity. In the presence of appropriate divalent cations such as Ca^2+^, HG chains form stable structures to enhance cell wall rigidity (Levesque-Tremblay *et al*., 2015). However, depending on the pH and the degree of methylesterification, both meHG and HG can be degraded by pectate lyases or polygalacturonases, thereby decreasing cell wall rigidity (Bonnin & Pelloux, 2020). We speculate that HG in the barley nucellus is resistant to hydrolysis and usually maintains cell wall rigidity, because cells in this region are deformed and elongated in HG-deficient *opm1 opm2* mutants (Fig. 3). Importantly, these changes in cell shape do not correlate with an obvious increase in the abundance of meHG epitopes. This suggests that PME substrates (i.e. meHG) are rapidly recycled in the ovule by other pectin modifying enzymes, both in wild-type and *opm1 opm2* mutants. An alternative possibility is that ovule meHG is masked by other polymers such as xylan or glucan (Chateigner-Boutin et al., 2014), the latter of which is present in the developing barley nucellus (Fig. S2).

It appears somewhat counterintuitive that in wild-type ovules, inner nucellar cells with a higher level of HG expand more than outer nucellar cells, even though HG is often implicated in limiting elasticity. However, it is important to note that recent models suggest HG is a structural component of the cell wall that supports adhesion between wall components and adjoining plant cells, thereby maintaining cell shape (reviewed in Du *et al*., 2022). Hence, the deposition of HG in inner nucellar cells may reflect an important structural role of these cells in resisting expansion of the female gametophyte, or in confining cell divisions to the outer nucellus and integuments. Consistent with the former possibility, we observed a wider ES in both *opm1 opm2* mutants and *pMADS31::PMEI-eGFP* lines. By contrast, the lack of cell proliferation in the inner nucellus does not appear to be directly dependent upon PME activity or HG accumulation, since cell division in the inner nucellus was not recovered in the *opm1 opm2* mutant (Fig 3).

Previous reports in multiple species provide some precedent for ovules exerting maternal influence on seed size by imposing physical restrictions, in particular via the seed coat, through modification of phytohormone pathways, and control of nutrient flow (Xie *et al*., 2015). In cereal species, physical constraints on seed size are also imposed by maternal floral organs (the lemma and palea), which define the limits for endosperm growth (reviewed in Li *et al*., 2018), and the pericarp, which coordinates early grain growth by expressing expansins to relax cell walls (Lizana *et al*., 2010). Ovary size also impacts grain size and/or weight (Guo *et al*., 2016), although this may relate to increased cell number rather than larger cells in wheat (Reale *et al*., 2017). Here, we identify another source of maternal control, which directly links ovule cell wall polysaccharide composition to grain development. Unlike the Arabidopsis ovule, where the nucellus is a small group of cells located around the ES and distal to the chalaza, the barley ovule incorporates a large volume of nucellar cells enclosing the ES. In a previous study of ovule development in 150 elite barley cultivars, nucellus area showed a strong positive correlation with ovule area, ovule length and ovule width (Wilkinson *et al*., 2019). Here we provide further support for this relationship by removing PME activity, a negative regulator of nucellus growth, which results in increased ovule length and width.

There are several possibilities to explain how the nucellus passes its maternal effects to grain growth. First, the nucellus affects the size of the ovule, and subsequently the ovary, which provides a congenital status in the grain precursor. In the case of the *opm1 opm2* mutant, ovaries expand more on the longitudinal axis than the transverse axis. Thus, the nucellus may fulfil a purely physical role in limiting longitudinal growth of the ovule and ovary, while the hull tissues (lemma and palea) limit lateral growth. Second, nucellus degradation is concomitant with endosperm cell number determination and ovary elongation during the first 3–4 days after pollination, which is critical for final grain shape (Ishimaru *et al*., 2003). In *opm1 opm2* ovules, the change in cell wall composition might accelerate nucellus degradation to affect early grain development. In future studies it will be important to understand how *OPM1* and *OPM2* expression is regulated, and to determine how OPM activity interacts with known pathways influencing seed size before and after fertilization.

## Acknowledgements

The authors wish to thank Chao Ma, Susan Johnson and Dr Gang Li for technical assistance and discussions. We also thank Dr Olena Kravchuk (Statistics for the Australian Grains Industry; SAGI) for significant support with statistical analysis. LCM experiments were undertaken with support from Dr Gwenda Mayo at Adelaide Microscopy. This research was supported by Australian Research Council grants to M.R.T (FT140100780 and DP180104092). KH would like to acknowledge funding from the Rural & Environment Science & Analytical Services Division of the Scottish Government.

## Author Contributions

XY and MRT designed the study. XY was responsible for phenotyping, molecular biology and microscopy experiments. LW and MA collected and generated LCM data and assisted in the immunolabelling methodology. NS analysed transcriptomic data. KH investigated allelic diversity in OPM1 and OPM2. XY and MRT wrote the manuscript. All authors contributed to the editing and reviewing of the manuscript.

