## Supporting Information for "Ovule cell wall composition is a maternal determinant of grain size in barley"

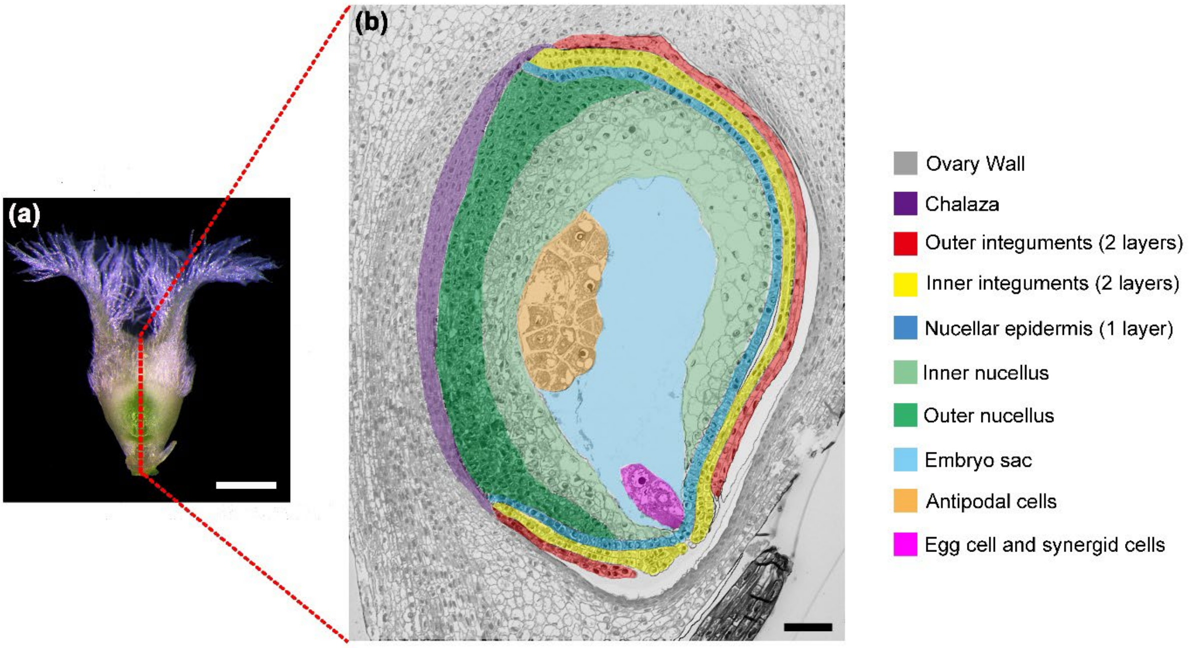


**Supporting Information Fig.S1: Anatomical structure of the barley ovule at anthesis (stage Ov10).**

**(a)** A barley pistil (cv. Golden Promise) at stage Ov10. The red dashed line indicates the position of the sagittal section in (b). Bar = 1mm.

**(b)** Semi-thin section of a barley ovule at anthesis, surrounded by ovary (carpel) tissues. Bar = 50 µm.

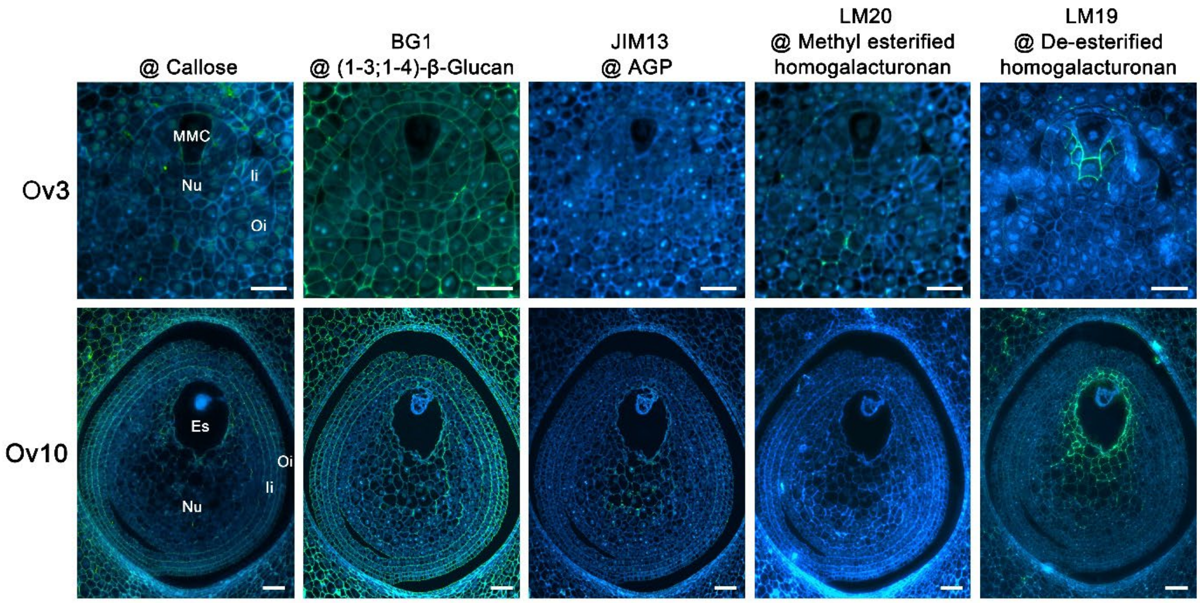


**Supporting Information Fig.S2: Immunohistochemical assay of various cell wall epitopes in the barley ovule.**

Callose, (1-3;1-4)-β-glucan, arabinogalactan proteins (AGP), methylesterified HG and de-esterified HG were detected by antibodies in wild-type ovules of two stages. Immunostaining is shown in green, Calcofluor White staining is shown in blue. Ov3, megaspore mother cell; Ov10, female gametophyte anthesis. OW, ovary wall; Oi, outer integument; Ii, inner integument; Nu, nucellus; ES, embryo sac. Bars = 25 µm.

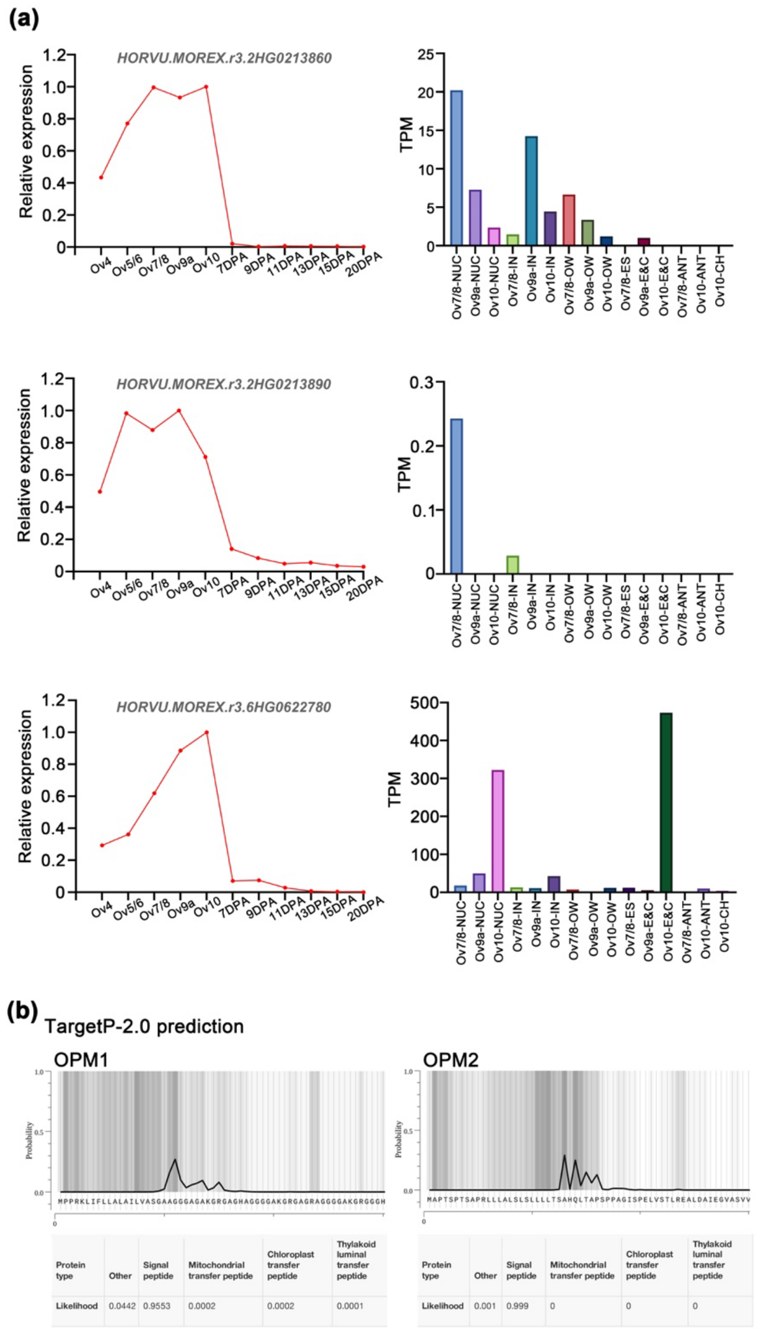


**Supporting Information Fig.S3: Molecular characteristics of OPM1 and OPM2.**

**(a)** Relative expression levels (left lane) of *OPM1* (HORVU.MOREX.r3.2HG0213860/HORVU.MOREX.r3.2HG0213890) and *OPM2* (HORVU.MOREX.r3.6HG0622780) in whole organs (pistil and caryopsis), as described by Matros et al. (2021), and transcripts per million (TPM) (right lane) of both genes in various tissues. Ov indicates pistil stages, DPA (days post anthesis) indicates caryopsis stages. Ov4, meiosis; Ov5/6, functional megaspore; Ov7/8, female gametophyte mitosis; Ov9a, mature female gametophyte; Ov9b, female gametophyte expansion; Ov10, female gametophyte anthesis.

**(b)** Predicted signal peptide (upper lane) and subcellular localisation (lower lane) of OPM1 and OPM2. Full length amino acids sequences were used for predictions in TargetP 2.0. The first 60 amino acids containing signal peptides are shown here.

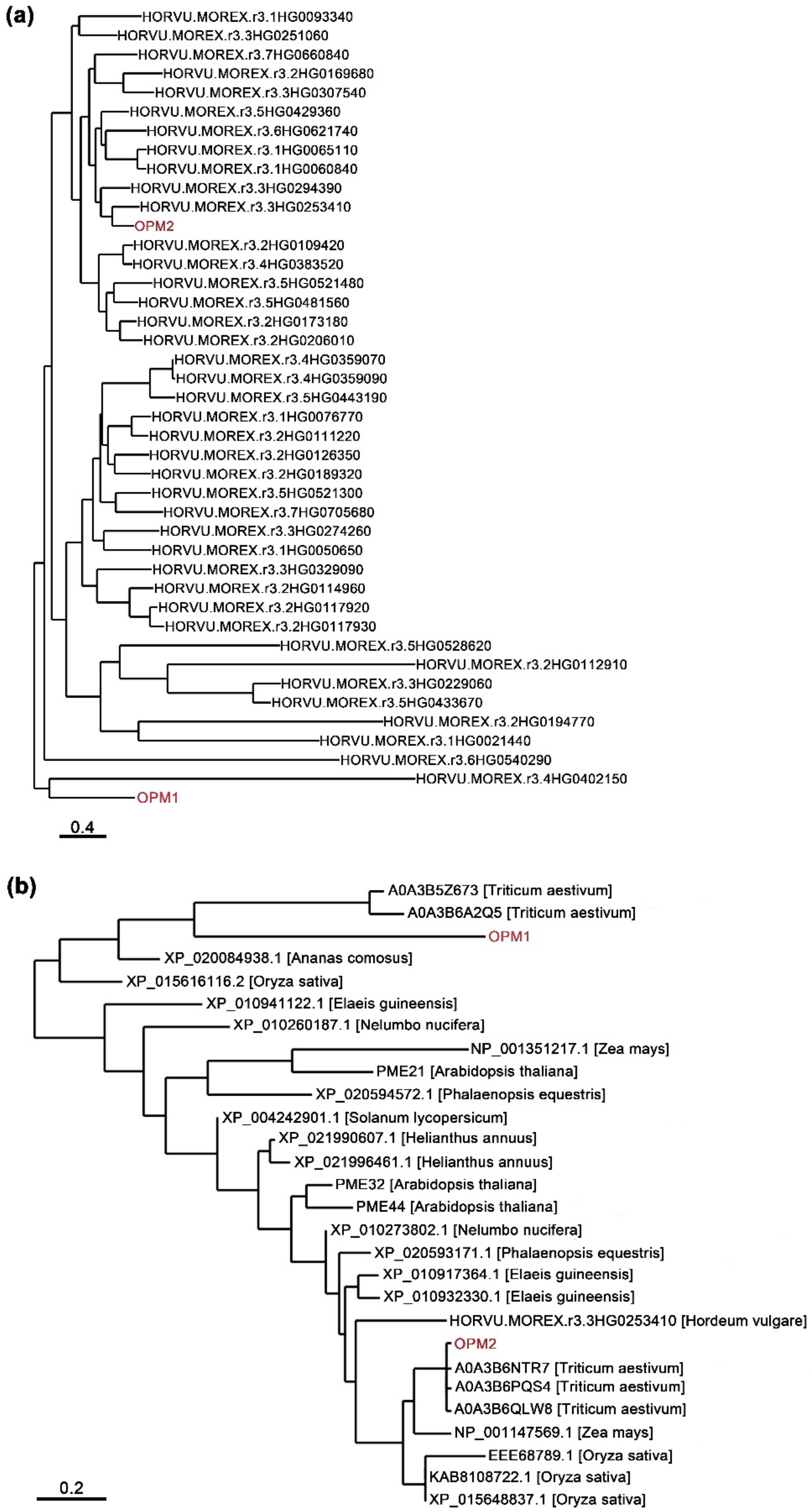


**Supporting Information Fig.S4: Phylogenetic analysis of OPM1 and OPM2 homologues.**

**(a)** Neighbour-joining tree of putative PMEs of barley showing OPM1 and OPM2 are not close homologues.

**(b)** Neighbour-joining tree showing homologues of OPM1 and OPM2 in higher plants.

**(a)**

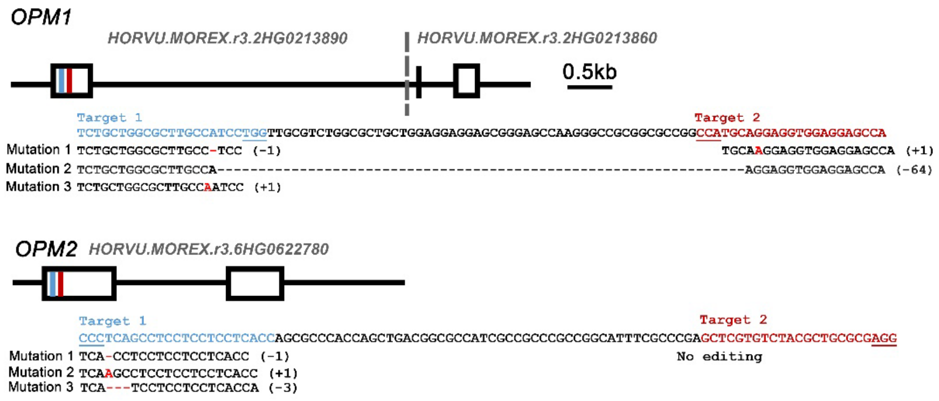


**(b)**

OPM1 MPPRKLIFLLALAILVASGAAGGGAGAKGRGAGHAGGGGAKGR

Mutation 1 MPPRKLIFLLALASWLRLALLEEEREPRAAAPAMQGGGGAKGR

Mutation 2 MPPRKLIFLLALAKEVEEPRAAAPAVQEVEEPRAAAAAMQEEVEHKASSPSPTALRTWS*

Mutation 3 MPPRKLIFLLALANPGCVWRCWRRSGSQGPRRRPCRRWRSQG

OPM2 MAPTSPTSAPRLLLALSLSLLLLTSAHQLTAPSPPAGISPELVSTL

Mutation 1 MAPTSPTSAPRLLLALSLASSSSPAPTS*

Mutation 2 MAPTSPTSAPRLLLALSLKPPPPHQRPPADGAIAARRHFARARVYA

Mutation 3 MAPTSPTSAPRLLLALSLI-LLLTSAHQLTAPSPPAGISPELVSTLREALDAIEGVAS

**(c)**

| | *opm1* | *opm2* | *opm1 opm2* |
|---|---|---|---|
| ***OPM1*** | Mutation 3 | WT | Mutation 1 |
| ***OPM2*** | WT | Mutation 2 | Mutation 1 |

**Supporting Information Fig. S5: Creation of *opm1 opm2* mutants using the CRISPR/Cas9 system.**

**(a)** Diagrams of gene structures for *OPM1* and *OPM2* and positions of two targets (blue and red). Targets were chosen close to the start of the coding sequence in an effort to generate null alleles. DNA sequences of independent mutations created by Cas9 are shown.

**(b)** The putative OPM1 and OPM2 amino acid sequences of wild type and mutants according to mutations identified from (a).

**(c)** Mutations in the mutants used in this study.

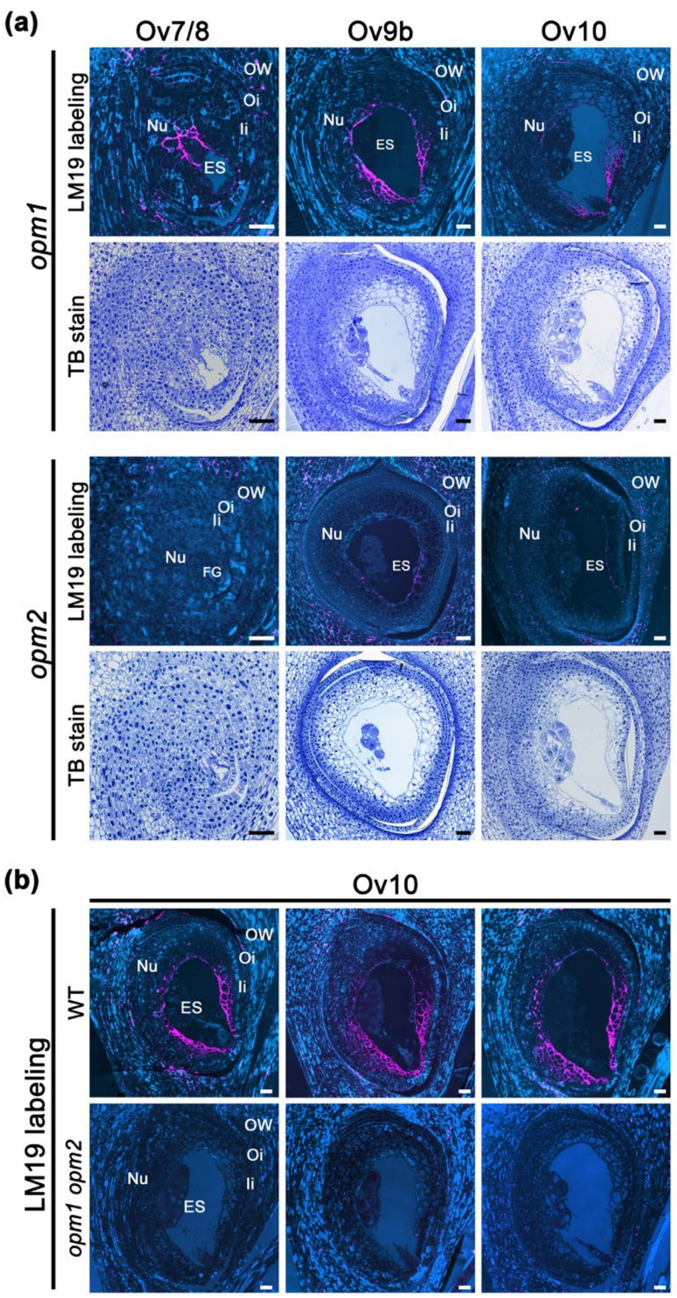


**Supporting Information Fig. S6: Phenotypic characterisation of ovules from single and double mutants.**

**(a)** Immunohistochemical assay of de-esterified HG (LM19) and cytological analysis of *opm1* and *opm2* ovules at different stages.

**(b)** Immunohistochemical assay of de-esterified HG (LM19) in three individual wild-type ovules (upper) and *opm1 opm2* ovules (lower), respectively.

Ov7/8, female gametophyte mitosis; Ov9b, female gametophyte expansion; Ov10, female gametophyte anthesis. OW, ovary wall; Oi, outer integument; Ii, inner integument; FG, female gametophyte; ES, embryo sac; TB, toluidine blue. Bars = 50 µm.

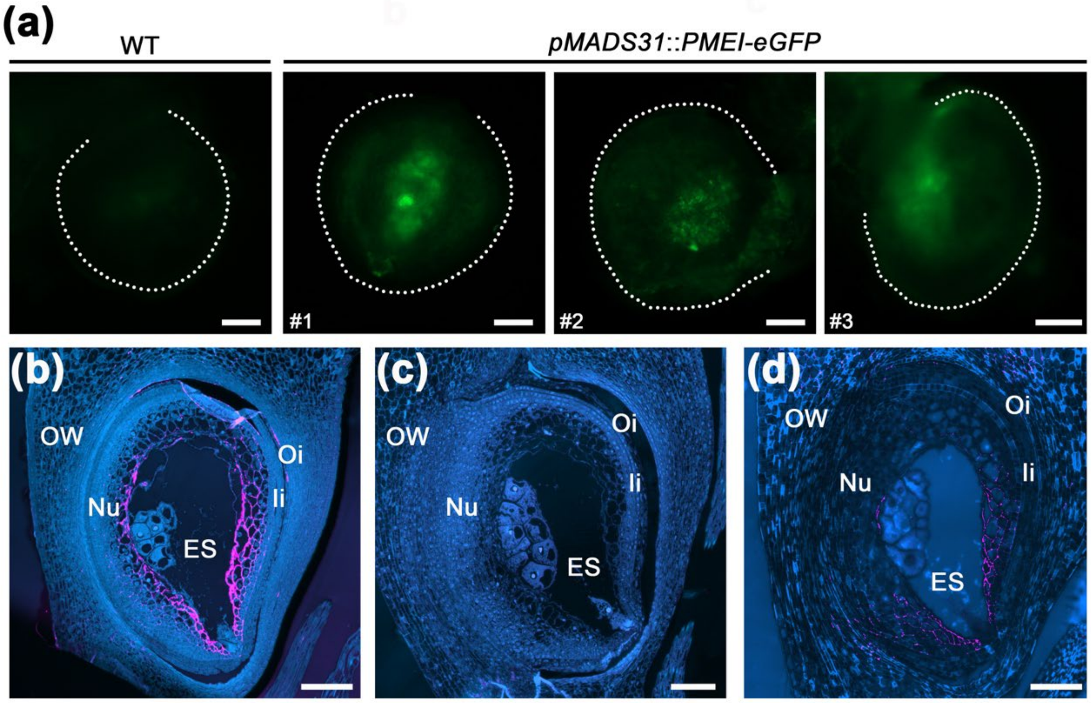


**Supporting Information Fig.S7: Characterisation of *pMADS31::PMEI-eGFP* transgenic plants.**

**(a)** Wild-type ovule and ovules expressing *pMADS31::PMEI-eGFP* from three independent transgenic lines.

**(b–d)** Immunohistochemical assay of de-esterified pectin (LM19) in wild-type (b), *opm1 opm2* (c) and *pMADS31::PMEI-eGFP* (d) ovules.

OW, ovary wall; Oi, outer integument; Ii, inner integument; ES, embryo sac. Dotted white line in (a) indicates the boundary of the ovule. Bars = 50 µm.

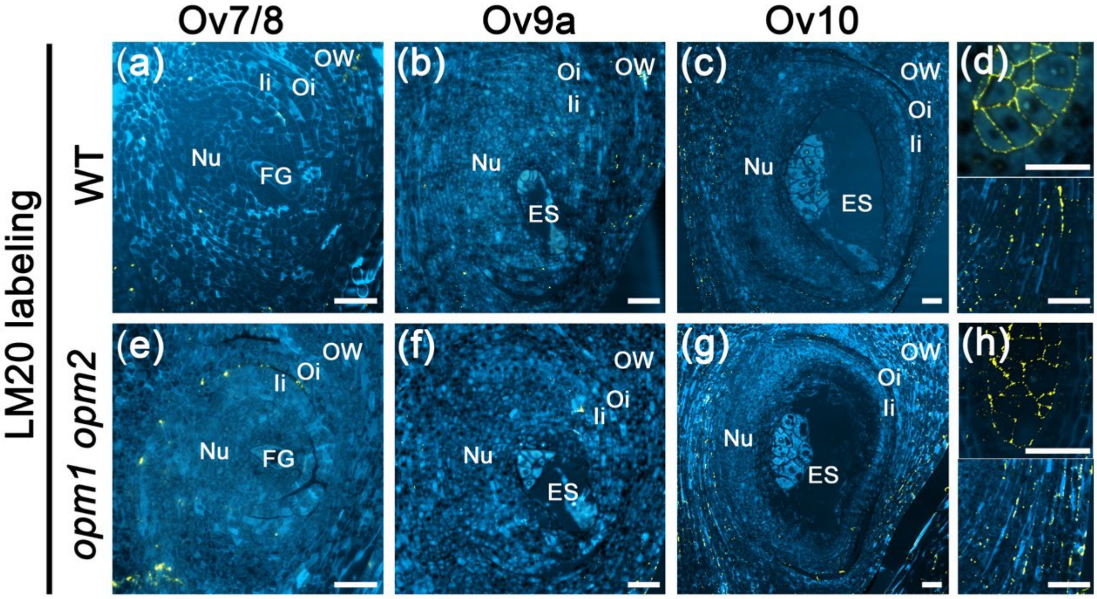


**Supporting Information Fig. S8: Immunohistochemical assay of meHG (LM20) in wild-type and *opm1 opm2* ovules at different stages.**

**(a–c)** LM20 labelling (yellow) in wild-type ovules.

**(d)** LM20 labelling in wild-type anthers (upper) and ovary wall (lower). Esterified pectin was detected in the walls of microspore mother cells and ovary cells (shown in yellow).

**(e–g)** LM20 labelling in *opm1 opm2* ovules.

**(h)** LM20 labelling in *opm1 opm2* anthers (upper) and ovary wall (lower). Esterified pectin localization was similar to that in wild type.

Ov7/8, female gametophyte mitosis; Ov9b, female gametophyte expansion; Ov10, female gametophyte anthesis. OW, ovary wall; Oi, outer integument; Ii, inner integument; FG, female gametophyte; ES, embryo sac. Bars = 50 µm.

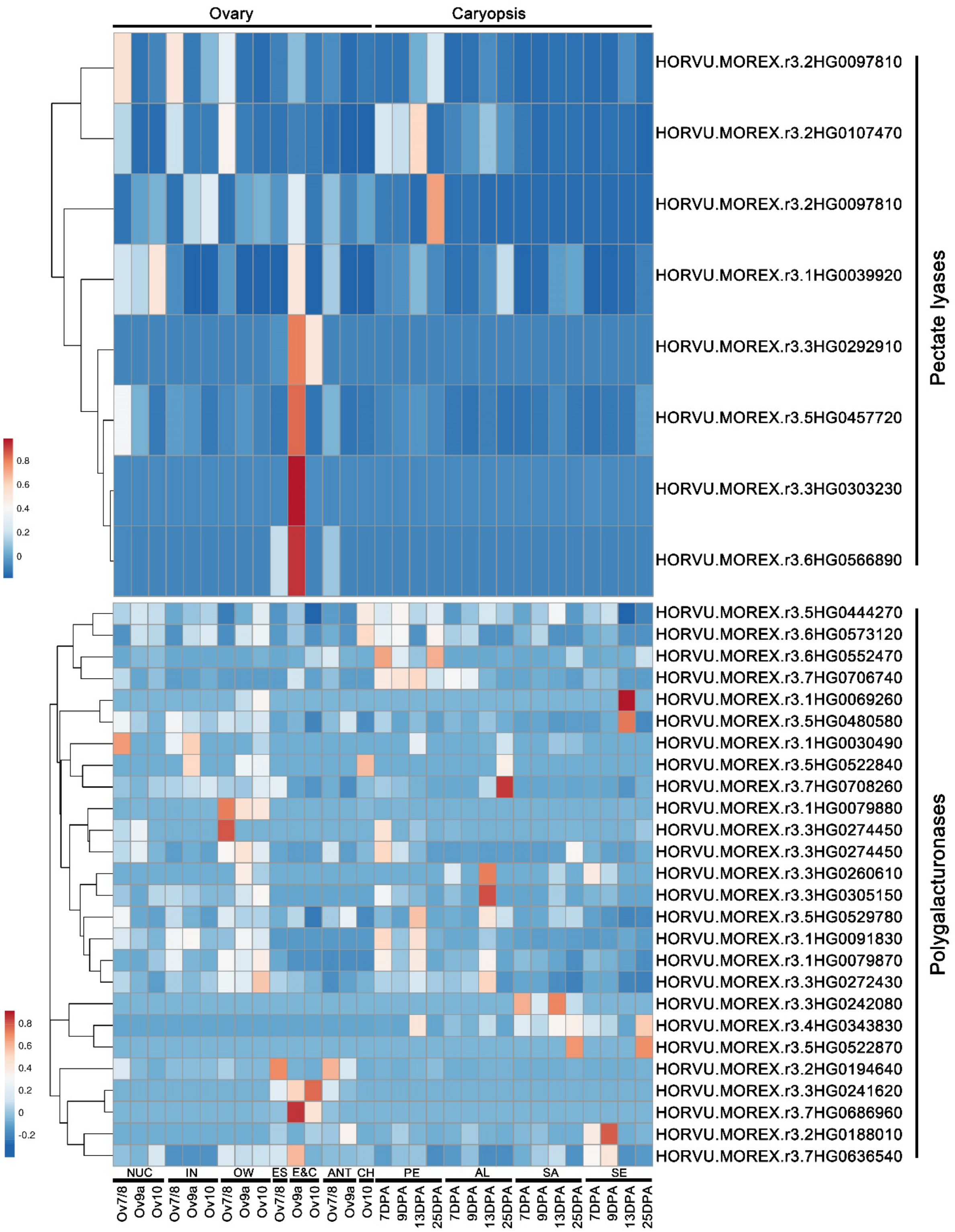


**Supporting Information Fig.S9: Heatmap visualization of the expression of pectin degradation genes in the ovary and caryopsis.**

Stages and tissues are indicated at the base of the figure: Ov7/8, female gametophyte mitosis; Ov9, female gametophyte maturity; Ov10, female gametophyte anthesis. DPA, days post anthesis. NUC, nucellus; IN, integuments; OW, ovary wall; ES, embryo sac; E&C, egg cell and central cell; ANT, antipodal cells; CH, chalaza; PE, pericarp; AL, aleurone; SA, sub-aleurone; SE, starchy endosperm.

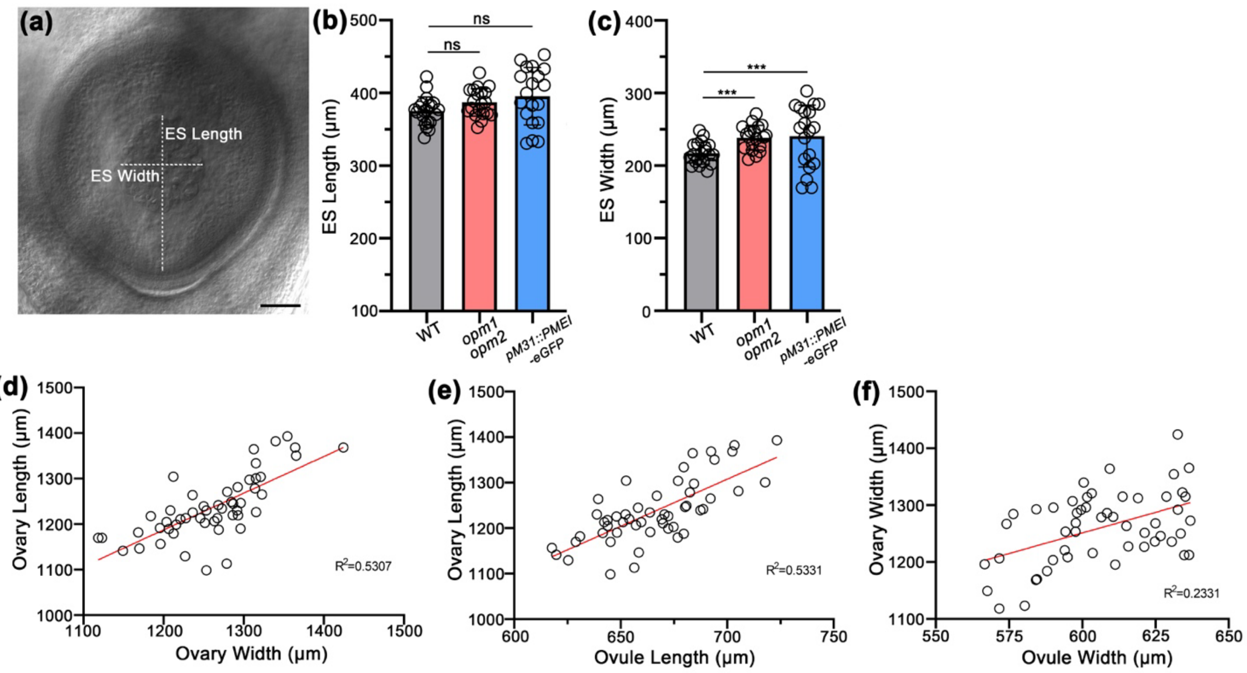


**Supporting Information Fig. S10: Embryo sac dimensions and correlations of ovule and ovary traits.**

**(a)** Embryo sac dimensions. Embryo sac length is the distance between the top of antipodal cell cluster and the micropyle. Embryo sac width indicates the width of the antipodal cells cluster. Bar = 50 μm.

**(b)** and **(c)** Measurements of embryo sac dimensions in wild-type, *opm1 opm2* and *pMADS31::PMEI-eGFP* ovules. All error bars represent SD, *n* = 20 (20 pistils from a pool of ~200 pistils from 6 plants). ANOVA test. ***, *P* ≤ 0.001; ns, not significant.

**(d)** Correlation between ovary width and ovary length. *n* = 60 pistils, $R^2$=0.5307.

**(e)** Correlation between ovule length and ovary length. *n* = 60 pistils, $R^2$=0.5331.

**(f)** Correlation between ovule width and ovary width. *n* = 60 pistils, $R^2$=0.2331.

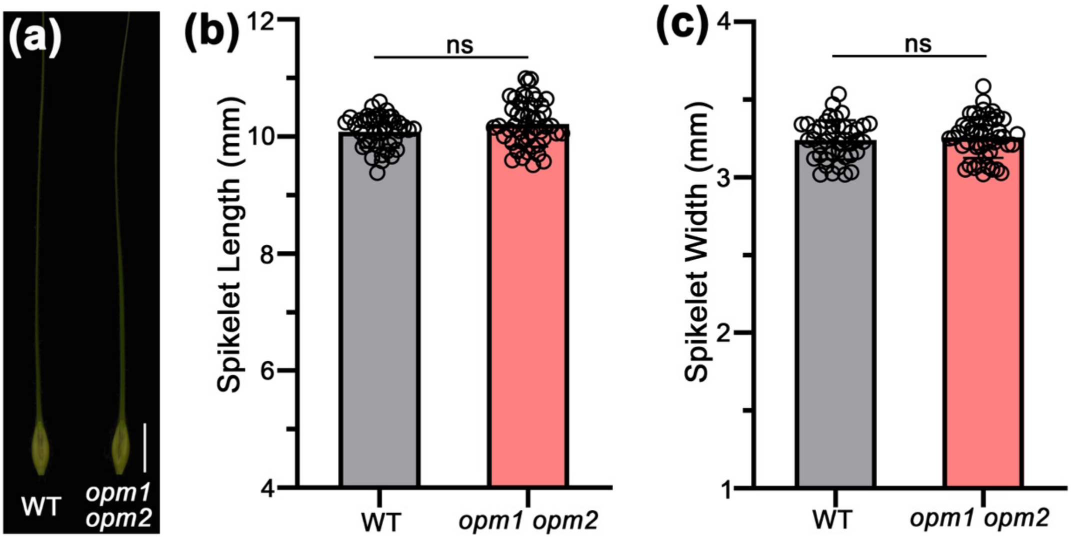


**Supporting Information Fig. S11: Measurements of spikelet size in wild type and *opm1 opm2* mutants.**

**(a)** Comparison of wild-type and *opm1 opm2* central spikelets. Bar = 1 cm.

**(b)** and **(c)** Comparisons of spikelet length (awns trimmed) and width in wild-type and *opm1 opm2* plants. All error bars represent SD, $n$ = 41 and 49 for WT and *opm1 opm2*, respectively (spikelets from 3 plants). Student $t$ test. ns, not significant. Further, an equivalence test (5% threshold) claims equivalence for both spikelet length and width.

(a)

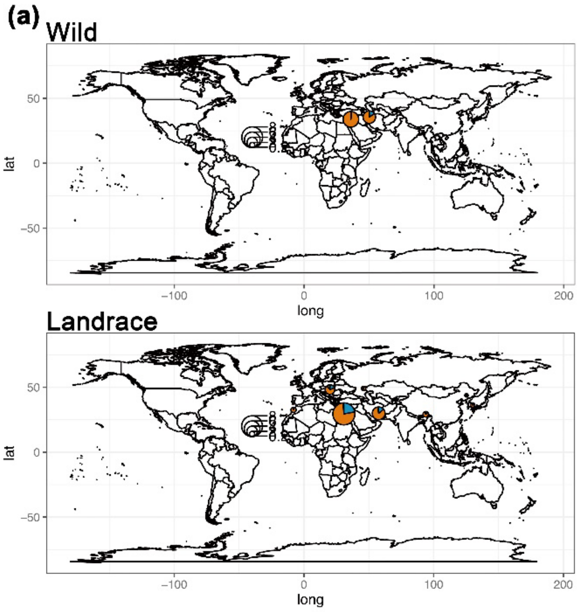


(b)

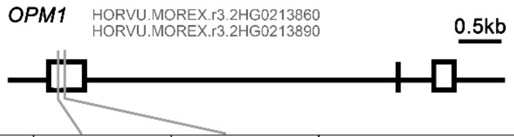


| Position | 755,729,635 | 755,729,724 | Subpopulation | |
|---|---|---|---|---|
| A.A.sub | A ➤ G<br>Asp ➤ Gly | C ➤ G<br>Asp ➤ His | Wild<br>(Total 109) | Landrace<br>(Total 209) |
| Hap A | A | C | 8 | 34 |
| Hap B | A | G | 101 | 164 |
| Hap C | G | G | 0 | 11 |

(c)

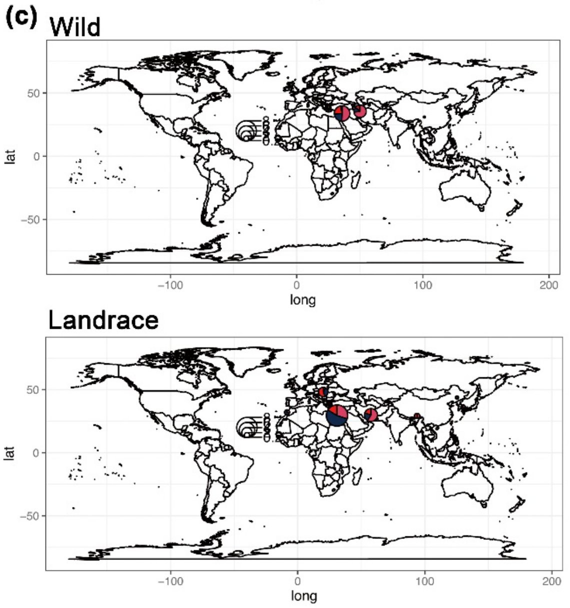


(d)

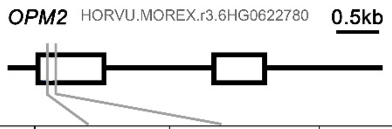


| Position | 553,211,304 | 553,211,344 | Subpopulation | |
|---|---|---|---|---|
| A.A.sub | C ➤ T<br>Val ➤ Val | G ➤ C<br>Ala ➤ Pro | Wild<br>(Total 105) | Landrace<br>(Total 210) |
| Hap A | C | G | 62 | 69 |
| Hap B | T | C | 20 | 90 |
| Hap C | T | G | 20 | 51 |
| Hap D | C | C | 3 | 0 |

**Supporting Information Fig. S12: SNP profiles of *OPM1* and *OPM2* in sequenced wild (*H. spontaneum*) and landrace barley accessions.**

The pie charts show the distribution of each haplotype (indicated at right and in the table) at each geographical location. *OPM1* contains two SNPs, 2H_755729635 (A/G) and 2H_755729724 (G/C) both of which change the amino acid sequence, either D109G or D139H respectively. These SNPs are not in linkage disequilibrium with each other and give rise to three haplotypes, although the rarer haplotype (represented by the nucleotides GG at these two SNPs) is not present in the *H. spontaneum* accessions and only 5% of the landraces. The most common haplotype was AG, (D109, D139) which is present in 93% of the *H. spontaneum* accessions and 78% of barley landraces. *OPM2* also contains two SNPs, 6H_553211304 (C/T) which is synonomous (V65V), and 6H_553211344 (G/C) which is non-synomous, resulting in A79P. This collection of *H. spontaneum* and barley landraces contained four haplotypes of *OPM2*, with the rarest haplotype in the *H. spontaneum* lines (CC, P79, 3% of *H. spontaneum* lines) being absent from the landrace accessions. The most common haplotype in the *H. spontaneum* accessions (CG, A79) was present in 59% of these lines, compared to 33% of the landrace accessions. The most common haplotype in the landraces (TC, P79) was observed in 43% of the landrace accessions include in this study compared to 19% of the *H. spontaneum* lines.

**Table S1. Sequences of primers used in this study.**

| Analysis | Name | Sequence (5' to 3') |
|---|---|---|
| **OPM1 genotype** | OP1-SF | CGCAATAAGTGGACTGGACAGCC |
| | OP1-SR | TGAGTGCGTCGGCAATGGAGGT |
| **OPM2 genotype** | OP2-SF | CAGTTCGAGTCGAGTGTTGGCGA |
| | OP2-SR | ACATGGACCACGAGAGCTCGTC |
| **CRISPR** | OP1T1U6-sgF | TCTGCTGGCGCTTGCCATCCGTTTTAGAGCTAGAAAT |
| | OP1T1U6a-R | GGATGGCAAGCGCCAGCAGACGGCAGCCAAGCCAGCA |
| | OP1T2U6-sgF | TGGCTCCTCCACCTCCTGCAGTTTTAGAGCTAGAAAT |
| | OP1T2U6b-R | TGCAGGAGGTGGAGGAGCCACAACACAAGCGGCAGC |
| | OP2T1U6-sgF | GTGAGGAGGAGGAGGCTGAGTTTTAGAGCTAGAAAT |
| | OP2T1U6c-R | TCAGCCTCCTCCTCCTCACCTGAGCCTCAGCGCAG |
| | OP2T2U3-sgF | GCTCGTGTCTACGCTGCGCGGTTTTAGAGCTAGAAAT |
| | Op2T2U3-R | CGCGCAGCGTAGACACGAGCTGCCACGGATCATCTGC |
| | U-F | CTCCGTTTTACCTGTGGAATCG |
| | gR-R | CGGAGGAAAATTCCATCCAC |
| | Pps-GGL | TTCAGAGGTCTCTCTCGACTAGTATGGAATCGGCAGCAAAGG |
| | Pgs-GG2 | AGCGTGGGTCTCGTCAGGGTCCATCCACTCCAAGCTC |
| | Pps-GG2 | TTCAGAGGTCTCTCTGACACTGGAATCGGCAGCAAAGG |
| | Pgs-GG3 | AGCGTGGGTCTCGTCTTCACTCCATCCACTCCAAGCTC |
| | Pps-GG3 | TTCAGAGGTCTCTAAGACTTTGGAATCGGCAGCAAAGG |
| | Pgs-GG4 | AGCGTGGGTCTCGAGTCCTTTCCATCCACTCCAAGCTC |
| | Pps-GG4 | TTCAGAGGTCTCTGACTACATGGAATCGGCAGCAAAGG |
| **qRT-PCR** | OPM1-RT-F2 | AAGCTTGTACGAAGGGGACG |
| | OPM1-RT-R2 | GTGCACTTGGGTCACTCACT |
| | OP2-RT-F | GGGAGTACATGAACACCGGG |
| | OP2-RT-R | CCACATGTTGCCCTCGATGA |
| | HvACTIN7-F | CGTGTTGGATTCTGGTGATG |
| | HvACTIN7-R | AGCCACATATGCGAGCTTCT |
| ***In situ*** | HvH4-ISH-AF | ATGTCTGGGCGTGGCAAGGG |
| **hybridization** | HvH4-ISH-AR | TAATACGACTCACTATAGGGTCAGCCGCCGAAGCCGT |
| | HvH4-ISH-SF | TAATACGACTCACTATAGGGATGTCTGGGCGTGGCAAGGG |
| | HvH4-ISH-SR | TCAGCCGCCGAAGCCGT |
| | OPM1-ISH-AF | ATGCGAAGGGCAGACGTTTA |
| | OPM1-ISH-AR | TAATACGACTCACTATAGGGATGCATGGAGTGGACGGATG |
| | OPM1-ISH-SF | TAATACGACTCACTATAGGGATGCGAAGGGCAGACGTTTA |
| | OPM1-ISH-SR | ATGCATGGAGTGGACGGATG |
| | OPM2U-ISH-AF | CAGCAACTTCACCGTCGCGCAGT |
| | OPM2U-ISH-AR | TAATACGACTCACTATAGGG GATAGCACGTACACGATTTGAAT |
| | OPM2U-ISH-SF | TAATACGACTCACTATAGGGCAGCAACTTCACCGTCGCGCAGT |
| | OPM2U-ISH-SR | GATAGCACGTACACGATTTGAAT |
| **Subcellular** | OPM1-GFP-F | GGACTCTTGACCATGGCCATGCCTCCGAGAAAGTTGATC |
| **localization** | OPM1-GFP-R | TGCTCACCATACTAGTTTGGGCCAAATCGCGCGCAT |

| | | |
|---|---|---|
| | OPM2-GFP-F | GGACTCTTGACCATGGCCATGGCGCCGACGTCTCCGAC |
| | OPM2-GFP-R | TGCTCACCATACTAGTAGACGCGAGCAGCCCGGACG |
| ***proMADS31::PMEI-eGFP*** | ProM31-F | GCAGGCATGCAAGCTTTGCCATCCTTGGTTTATGTCACTG |
| | ProM31-PMEI-R | GCCAACGGTACGCGCCATCGCTCCAGCCCACTGGATCTCT |
| | PMEI-GFP-F | ATCCAGTGGGCTGGAGCGATGGCGCGTACCGTTGGCTGCC |
| | PMEI-GFP-R | TGCTCACCATACTAGTTGGCGGCGCCGTTTGGTTGGCC |
